# The functional organization of retinal input to the mouse superior colliculus

**DOI:** 10.64898/2026.04.15.718783

**Authors:** Yongrong Qiu, Lisa Schmors, Na Zhou, Mels Akhmetali, Dominic Gonschorek, Cameron Smith, Anton Sumser, Marie Vallens, Cathryn R. Cadwell, Fabrizio Gabbiani, Maximilian Joesch, Andreas Tolias, Philipp Berens, Thomas Euler, Fabian Sinz, Jacob Reimer, Katrin Franke

## Abstract

The superior colliculus integrates retinal input to drive rapid, adaptive visual behavior, yet how the functional diversity of retinal ganglion cell types is represented in superior colliculus remains poorly understood. Using chronic two-photon calcium imaging of retinal ganglion cell axonal boutons in awake mice, we recorded over 200,000 boutons across superficial superior colliculus layers — a scale that enabled systematic comparison with large-scale *ex vivo* retinal datasets. This revealed that the superior colliculus receives a near-complete sampling of retinal ganglion cell functional diversity. Functionally distinct response types were organized in systematic laminar gradients: not only response properties such as direction selectivity and contrast suppression, but retinal response types themselves varied systematically with depth. To probe how this organized input encodes natural scenes, we trained a “digital twin” deep network model on natural movie responses and validated its generalization to parametric stimuli, including cell type identification. Leveraging this model to generate predicted responses to looming stimuli, we identify a discrete subset of retinal response types tuned for collision detection at low angular thresholds — a specialization embedded within a broader, non-specialized retinal population. The digital twin is made publicly available as a community resource. Together, these findings provide a comprehensive functional map of retinal drive to the superior colliculus and an *in silico* platform for linking retinal cell types to behaviorally relevant superior colliculus computations.

## Introduction

The superior colliculus (SC), the mammalian homolog of the optic tectum (reviewed in Isa et al., 2021; Farrow et al., 2019), orchestrates rapid behavioral responses to visual threats and opportunities—from predator avoidance (e.g. Shang et al., 2015; Evans et al., 2018; Baier et al., 2025) and prey capture (e.g. Hoy et al., 2016, 2019) to orienting movements (e.g. Hafed, 2018; Bogadhi and Hafed, 2023). In mice, these diverse behaviors are driven by a feedforward projection from the majority of retinal ganglion cells (RGCs) (Ellis et al., 2016). Because the retinal output consists of over 40 well-characterized cell types (Baden et al., 2016; Bae et al., 2018; Goetz et al., 2022) and SC neurons map onto defined motor outputs (e.g. Hoy et al., 2019), the retino-collicular pathway offers an excellent model system for dissecting sensorimotor processing. While individual retino-collicular circuits have been characterized for specific behaviors (e.g. Hoy et al., 2019; Reinhard et al., 2019; Kühn et al., 2025), how the full diversity of parallel retinal channels is functionally organized within the SC remains unknown. Yet this organization is critical for understanding how SC neurons transform retinal input into coherent motor commands — as suggested by their demonstrably reduced dimensionality and novel feature selectivity relative to their inputs (Lee et al., 2020; Li and Meister, 2023a). Here, we address this gap by characterizing the functional organization of retinal input to the SC in the awake animal, thereby defining the input types that form the building blocks of downstream collicular integration and computation.

Several lines of work have begun to illuminate the organization of retinal input to the SC, but key questions remain. Anatomical studies have shown that different RGC types terminate at distinct depths within the superior colliculus (Reinhard et al., 2019; Huberman et al., 2008; Hong et al., 2011). Electrophysiological recordings have sampled retinal inputs with biases toward particular RGC classes (Sibille et al., 2022), while calcium imaging of RGC axon terminals has primarily addressed saliency encoding (Borges et al., 2025) or behavioral modulation (Schröder et al., 2020) rather than defining input cell types. Consequently, it remains unclear how the full diversity of retinal signals is organized at SC input. Furthermore, effectively linking these inputs to behavioral outcomes requires computational tools that can generalize beyond standard parametric stimuli to scale across the full diversity of behaviorally relevant inputs.

Here, we address these challenges by combining chronic two-photon imaging of RGC axonal boutons in the SC of awake mice with data-driven computational modeling, so-called functional “digital twin” models that predict the activity of individual neurons or boutons given the visual stimulus (Klindt et al., 2017; McIntosh et al., 2016; Batty et al., 2017; Sinz et al., 2018; Lurz et al., 2020a; Franke et al., 2022; Turishcheva et al., 2024; Wang et al., 2025; Willeke et al., 2025). Using a custom cranial window that enabled stable recordings over extended periods, we directly measured the activity of large populations of RGC inputs across SC depth, allowing us to define functional retinal response types and their laminar organization. By relating these measurements to an available large-scale *ex vivo* RGC dataset (Gonschorek et al., 2025), we showed that retinal functional diversity is largely preserved at the level of SC afferents and organized across collicular depth in a feature- and response type-specific manner.

To move beyond descriptive mapping using parametric stimuli and enable systematic interrogation of how RGC input supports behaviorally relevant computations in the SC, we trained a functional digital twin model on responses to natural movies. We showed that this model generalizes from natural movies to parametric out-of-distribution stimuli and captures known organizational properties of SC input, including retinotopy, recording depth, and retinal cell types identified based on (Baden et al., 2016). We further demonstrate how the model can be used to generate testable predictions about behavioral relevance by applying it to looming stimuli, an established paradigm mimicking approaching predators — narrowing the candidate retinal response types likely to underlie SC-mediated predator avoidance to a small subset. We will make the digital twin freely available, enabling reproducible and extensible *in silico* analyses of retinal drive to the SC under naturalistic and ethologically relevant visual conditions.

### Results

#### Two-photon imaging of retinal axons in mouse superior colliculus: Stable responses to parametric and natural movies

To measure retinal input to the superior colliculus (SC), we used intraocular GCaMP8m delivery and chronic cranial window implantation (Fig. 1a). Specifically, GCaMP8m was expressed under a pan-neuronal promoter, resulting in labeling of essentially all retinal ganglion cells (RGCs) and their axons projecting to the SC. To access these inputs in the SC, we implanted a custom 3D-printed cranial window designed to accommodate the overlying cortex. The cortical tissue was carefully displaced laterally but not aspirated, allowing stable optical access to a large fraction of the SC. Post hoc histological analysis in a subset of animals confirmed accurate placement of the window and revealed no detectable immune response, as assessed by Iba1 staining ((Fig. 1b); Holtmaat et al., 2009). The preparation was stable, supporting repeated imaging sessions for up to six months.

**Fig. 1.**
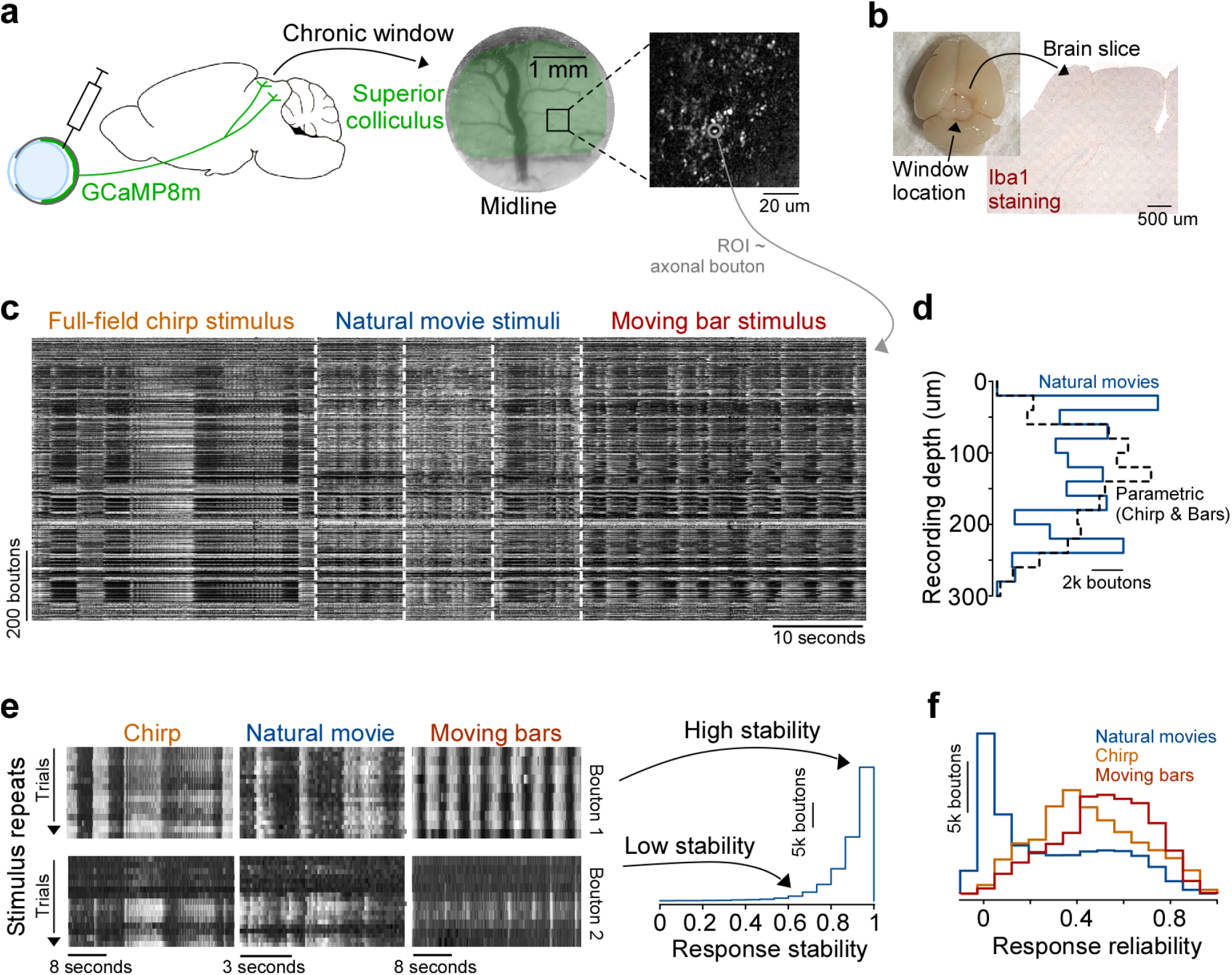
Two-photon imaging of retinal axons in mouse superior colliculus: stable responses to parametric and natural movies. **a**, Schematic of intraocular injection and chronic window implantation over the mouse superior colliculus (SC) for *in vivo* two-photon calcium imaging. AAVs expressing GCaMP8m under a pan-neuronal promoter was injected into the vitreous of anesthetized mice, leading to expression in retinal ganglion cells projecting their axons to the SC. A 3-mm cranial window was then implanted over the SC, with cortical tissue carefully displaced. Imaging was performed in awake, head-fixed mice to record visually evoked calcium activity from individual axonal boutons. **b**, Photograph of an explanted brain after perfusion, showing the location of the implanted window, alongside a histological section stained for Iba1 to assess immune response. **c**, Example single-trial calcium responses from 650 axonal boutons in one session, evoked by three stimulus types: a full-field “chirp” stimulus with contrast and frequency modulations (orange), natural movie clips (blue), and a bright bar moving in eight directions (red). **d**, Depth distribution of recorded boutons, referenced to the SC surface (0 *µ*m). Distributions are shown separately for boutons recorded with natural movie versus parametric stimuli. **e**, Example bouton responses illustrating trial-to-trial stability for each stimulus type, where stability is quantified as the slope of leave-one-out correlations across successive trials (see Methods). Top row: bouton with stable responses (near-zero slope) across a ∼2-hour recording. Bottom row: bouton with unstable responses (significantly non-zero slope). Histogram (right) shows the distribution of stability slopes for natural movie responses across all boutons; boutons exceeding the drift threshold (dashed line) were excluded from further analysis. **f**, Distribution of response reliability across boutons for each stimulus type. Reliability is defined as the mean leave-one-out correlation across all trial repetitions, collapsing across trial order to yield a single scalar measure of response consistency. Values range from 1 (identical responses across repeats) to 0 (uncorrelated responses, i.e. noise).

Using this preparation, we imaged retinal axonal boutons in up to five optical planes distributed in depth across the superficial right SC in awake, head-fixed mice (n = 201,352 boutons, n = 42 recordings, n = 3 mice). We presented three stimulus classes (Fig. 1c): a full-field “chirp” with temporally modulated contrast and frequency, a bright bar moving in eight directions, and natural movie clips. In each experiment, animals were shown either parametric stimuli alone (chirps and moving bars presented within the same recording) or natural movie trials interleaved randomly with parametric trials. Chirp and moving bar stimuli were chosen because they are classical paradigms for clustering functional retinal cell types (Baden et al., 2016), whereas natural movies provided ecologically relevant visual input suitable for training neural predictive models of RGC axonal boutons (for details on the movie stimuli, see Wang et al., 2025; Turishcheva et al., 2023). Boutons were recorded from the SC surface down to 300 µm in depth, with similar distributions across parametric and natural movie recordings (Fig. 1d).

To assess the stability of bouton responses over the recording session, we computed leave-one-out correlations across stimulus repetitions: for each trial, we correlated that trial’s response with the mean response computed across all remaining trials, thereby obtaining a per-trial estimate of response reliability. We then quantified monotonic drift by regressing these per-trial correlations against trial order. Boutons with an absolute slope smaller than a set threshold (0.2) were classified as monotonically drifting and, thus, were excluded (Fig. 1e). In addition, we quantified response reliability as the average leave-one-out correlation across trials. Parametric stimuli (chirps and moving bars) evoked more reliable responses than natural movies (Fig. 1f), consistent with their design to strongly drive retinal inputs with high-contrast features. For all further analyses, we included boutons only if they met stability and/or reliability criteria (see Methods for details).

### A systematic functional characterization of the retinal input to the SC

Having established a stable preparation for long-term imaging of retinal axonal boutons in the SC, we next asked how many functionally distinct response types of retinal input could be resolved with this approach. We began by focusing on responses to parametric visual stimuli—chirps and moving bars—because they are well established for dissecting and classifying RGC types based on temporal dynamics, contrast sensitivity, and local motion selectivity (Baden et al., 2016; Goetz et al., 2022).

We identified functionally distinct retinal response types using Gaussian mixture model (GMM) clustering on temporal features extracted by sparse principal component analysis (sPCA; see Methods and Fig. 1). Our analysis revealed 50 functionally distinct clusters (Suppl. Fig. 1), which we refer to as “response types” rather than cell types because they are defined solely by bouton response dynamics to the presented visual stimuli, without independent anatomical, molecular, or connectivity validation. We arranged the types according to a hierarchical tree based on their functional similarity (Fig. 2a) and assigned descriptive labels based on canonical response characteristics (e.g., On, Off, On-Off, suppressed-by-contrast).

**Fig. 2.**
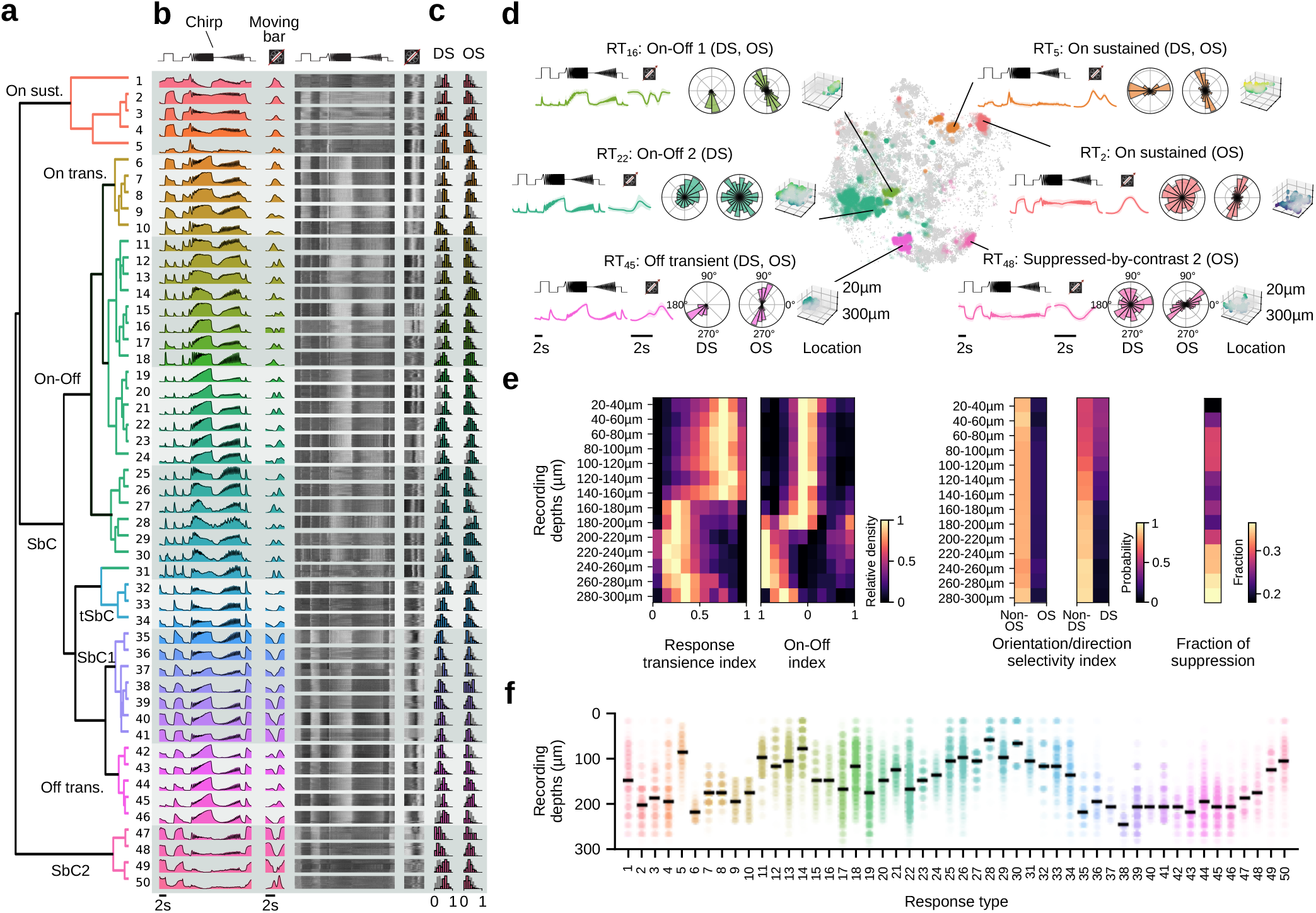
Retinal inputs to the SC form functionally distinct, laminar-organized response types. **a**, Cluster-dendrogram with response types indicated: *n*_ON_ = 10, *n*_OFF_ = 5, *n*_ON-OFF_ = 21, *n*_Sbc_ = 14 types. **b**, Cluster-mean Ca^2+^ (left) and single trial responses (right) to the two parametric stimuli, a full field chirp stimulus and local motion bar stimuli. **c**, Direction selectivity index and orientation selectivity index. Background histograms demarcate all retinal axon boutons. **d**, Two dimensional embedding of all 70,000 retinal axon bouton responses using the contrastive learning dimensionality reduction algorithm TRACE with example clusters indicated. Each example shows (from left to right) the mean response to ‘chirp’ and moving bar stimulus with standard deviation, the direction selectivity, the orientation selectivity in polar plots, and the anatomical location for the respective cluster. **e** Response properties across depths with (from left to right) the response transience index, the On–Off index, the orientation and direction selectivity, and the fraction of responses showing a modulation by contrast. **f**, Depth distribution across response types. Black bars indicate mean across boutons assigned to this type.

Our framework yielded a diverse set of retinal response types with distinct functional properties. Responses to the step component of the chirp stimulus comprised On, Off, On-Off, and Suppressed-by-Contrast (SbC) groups with both sustained and transient temporal dynamics (Fig. 2a-e). Responses were also highly modulated by contrast and frequency components of the chirp stimulus. Retinal inputs showed diverse selectivity to moving bar stimuli, with some showing direction and orientation selectivity (quantified by direction and orientation selectivity indices, DS and OS; Fig. 2d and Suppl. Fig. 2). To confirm that these functional distinctions were not an artifact of the clustering procedure, we examined whether they were preserved in a clustering-independent low-dimensional embedding of response dynamics. Two-dimensional TRACE embeddings (Schmors et al., 2025) recapitulated the organization revealed by GMM clustering, with response types forming well-separated manifolds according to direction selectivity, orientation selectivity, and depth (Fig. 2d; Suppl. Fig. 3). This convergence of cluster-based and embedding-based analyses provides independent support for the functional distinctiveness of the identified response types.

Importantly, we found that functional properties, such as direction selectivity, orientation selectivity, response transience, and contrast sensitivity (quantified by the suppression index, SI) varied systematically along the depth axis of the SC (Fig. 2e). Direction selective responses were predominantly found superficially (0-150*µ*m) but rare in deeper regions of the SC (150-300*µ*m; Fig. 2e, middle), in line with prior work (Inayat et al., 2015). Similarly, the transience index, which describes response kinetics to full-field light steps, showed a sharp transition from highly transient responses superficially to sustained responses in deeper regions (Fig. 2e, left). The On–Off index also varied with depth: deeper SC regions >180*µ*m deep showed strong selectivity for either light onset or offset, while superficial regions (0-180*µ*m) had an On-Off index near zero (Fig. 2e, left). Additionally, the contrast suppression, quantified by the suppression index (SI), changed across depth, with contrast-suppressed responses predominant at depths of 200-300*µ*m (Fig. 2e, right).

Notably, this laminar organization was also evident at the level of individual response types: although depth was not used as a feature during clustering, most response types projected to restricted depth ranges within the SC (Fig. 2f). The fact that functionally defined types segregate along the depth axis provides independent support for the biological validity of the clustering, and suggests that laminar position is a reliable correlate of retinal input identity in the SC.

Taken together, our data reveal substantial functional diversity in the retinal input to the SC, with functional properties systematically organized along the depth axis and functionally related inputs clustering within specific laminar zones.

### Functional diversity of *ex vivo* retinal output matches that of SC retinal input

To determine whether the SC receives the full range of functional diversity present in the retinal output (Baden et al., 2016; Goetz et al., 2022), we aligned our *in vivo* retinal axonal bouton recordings with a large-scale *ex vivo* dataset of RGC responses using batch-integration techniques (Gonschorek et al., 2025; Ganin et al., 2016). We utilized this reference dataset comprising >26,000 RGCs recorded from the ganglion cell layer of the mouse retina, with characterized responses to full-field chirp and moving bar stimuli.

Aligning these datasets, one recorded *ex vivo* from the retina and the other *in vivo* from awake, behaving mice, presented significant challenges due to domain-specific differences, including distinct calcium indicators (*ex vivo* with Oregon Green BAPTA-1 (OGB-1) vs. *in vivo* with GCaMP8m), experimental preparations and potential response modulation through synaptic input to RGC terminals *in vivo*. Consistent with these differences, low-dimensional embeddings of the raw response features revealed a strong separation between datasets (Fig. 3a), indicating that domain identity dominated the representational space.

**Fig. 3.**
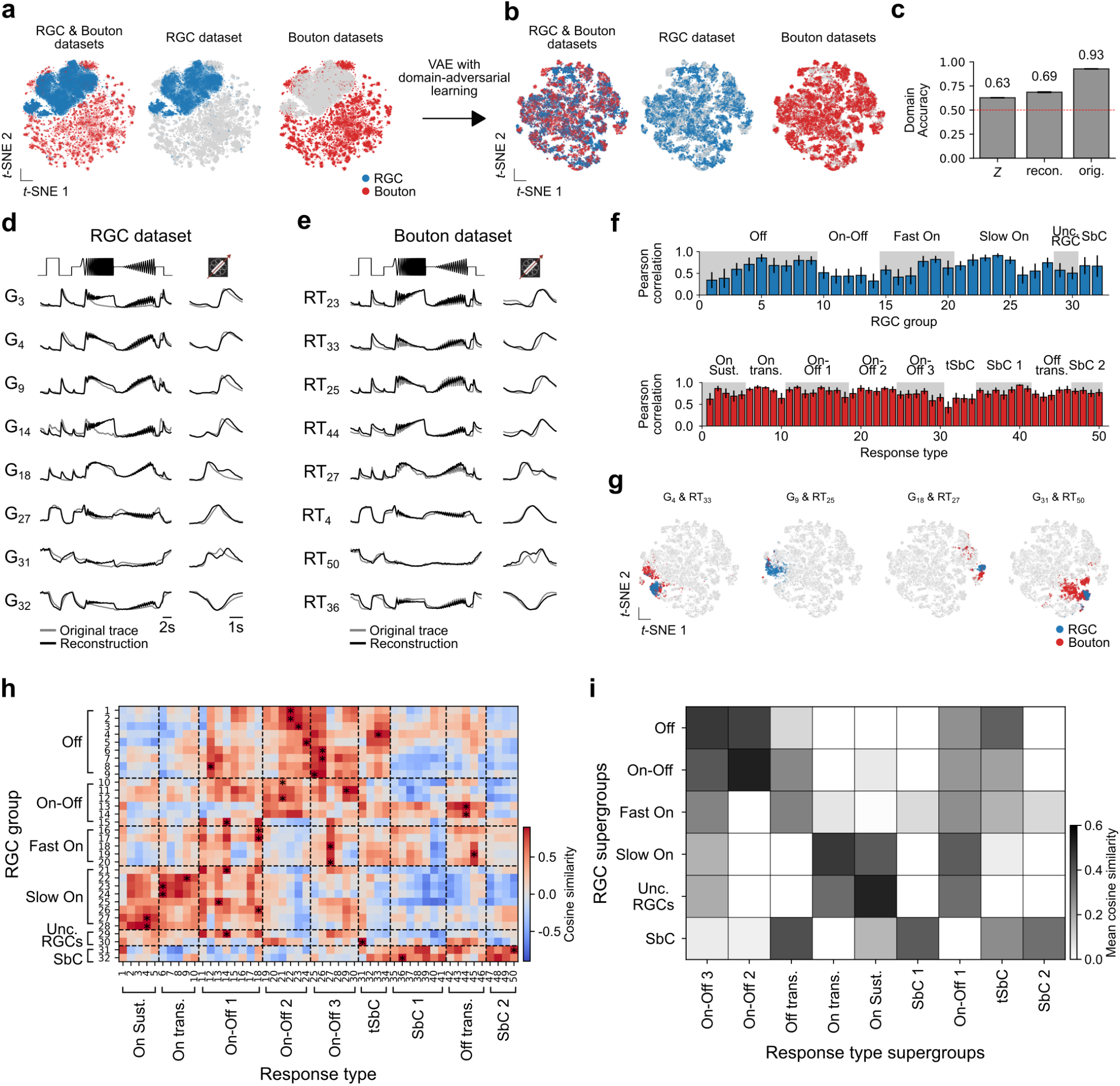
Matching *ex-vivo* RGC groups with in-vivo bouton response types. **a**, *t*-SNE visualization of latent representations from retinal ganglion cell (RGC, blue) and retinal bouton (red) datasets. Left: joint embedding without domain alignment shows separation by dataset. Middle and right: embeddings of each dataset separately. **b**, *t*-SNE visualization after training a variational autoencoder (VAE) with domain-adversarial learning. Joint embedding (left) shows strong mixing of RGC and bouton response type samples, indicating domain alignment, while dataset-specific embeddings (middle, right) preserve internal structure. **c**, Domain classification accuracy from latent features (*Z*), reconstructed traces, and original traces, used as a measure of domain invariance. A classifier trained to predict the recording domain (e.g., animal or session identity) from the representation should perform at chance level if the representation is fully domain-invariant; accuracy above chance indicates residual domain-specific information. Dashed line marks chance level; error bars represent cross-validation variability. **d**, Example RGC group responses to the full-field ‘chirp’ (left) and moving bars (right). For each group, mean original traces (gray) and reconstructions (black) are shown. **e**, Example bouton response types shown as in (d). **f**, Reconstruction performance quantified as Pearson correlation between original and reconstructed responses for all RGC groups (top, blue) and bouton response types (bottom, red). Bars indicate mean correlation per group/cluster. Functional supergroups are indicated above. **g**, Example correspondences between selected RGC groups and bouton response types visualized in the shared latent space using *t*-SNE. Highlighted points show cells from matched groups/clusters overlaid on the full embedding. **h**, Cosine similarity matrix between mean response profiles of RGC groups (rows) and bouton response types (columns). Dashed lines indicate functional supergroup boundaries; asterisks mark highest similarities per RGC group. **i**, Mean cosine similarity aggregated at the level of RGC and bouton response type functional supergroups, summarizing cross-structure correspondence.

To bridge this gap, we employed a variational autoencoder with adversarial training to reduce dataset-specific domain information while preserving biologically meaningful signals. In this framework, the model learns a shared latent representation *Z* from which reconstruction of the original responses remains accurate, while an adversarial classifier is trained to minimize the decodability of the recording domain (*ex vivo* vs. *in vivo*) from *Z*. This approach follows the general paradigm of domain-adversarial representation learning (e.g. Ganin et al., 2016), which has been widely applied across fields including medical imaging (e.g. Kamnitsas et al., 2017) and genomics (e.g. Dincer et al., 2020), as well as systems neuroscience (e.g. Gonschorek et al., 2021).

After training, latent representations from the RGC and bouton datasets showed strong mixing, indicating successful domain alignment (Fig. 3b). Consistent with this, a logistic regression domain classifier trained on the latent representation performed near chance level, indicating reduced domain-predictive information compared to both the original feature space and reconstruction-only embeddings. (Fig. 3c). Importantly, stimulus-evoked response structure was preserved: reconstructed traces closely matched the original responses for representative RGC groups and bouton response types across both chirp and moving bar stimuli (Fig. 3d–f). For the moving bar, directional information was discarded and responses were averaged across directions to focus on shared temporal response structure. Overall, reconstruction quality was higher for bouton responses than for RGC responses, as quantified by the Pearson correlation between original and reconstructed traces (Fig. 3f). While bouton reconstructions showed consistently high correlations (Fig. 3f, bottom), certain RGC groups (e.g. Off and On–Off) exhibited lower reconstruction performance (Fig. 3f, top), likely reflecting differences in signal-to-noise ratio and group-definition procedures between the two datasets.

We next quantified functional correspondence between RGC groups and bouton response types. For this analysis, we computed mean latent representations (“group centroids”; examples in Fig. 3g) for each RGC group and each bouton response type, and measured their pairwise cosine similarity to obtain a cross-dataset similarity matrix (Fig. 3h). For each RGC group, at least one bouton response type exhibited high latent similarity. However, substantial similarity was also observed among different RGC groups, indicating that several retinal types share closely related latent representations. Thus, cross-structure correspondences likely reflect shared functional motifs rather than strict one-to-one type identity (Fig. 3h, asterisks). Consistent with this, multiple RGC groups often mapped to the same bouton response type, reflecting shared response structure.

To summarize these relationships across broader functional categories, we aggregated similarities across predefined RGC and bouton functional supergroups (e.g. Transient On, Sustained Off). This analysis revealed structured correspondences between major response classes, with several supergroups exhibiting elevated cross-structure similarity (Fig. 3i). Although these mappings were not strictly one-to-one, they demonstrate that core functional motifs of retinal output are preserved in retinal axon boutons across SC layers. This preservation suggests that local SC circuits minimally transform the functional signatures of incoming retinal signals, at least at the level of these canonical response properties.

Together, these analyses indicate that, after domain alignment, retinal axon bouton responses can be systematically related to RGC functional groups and capture a broad subset of retinal functional diversity. This supports the view that much of the functional organization present in the retina is preserved in its major retino-recipient target, while potentially being selectively weighted toward specific response classes.

### A high-performance functional digital twin of retinal input to the SC across animals

Our analyses so far revealed a rich, laminar organization of retinal input to the SC and demonstrated that its functional diversity closely mirrors that of RGCs characterized *ex vivo*. The next challenge is to understand how this diverse retinal code represents behaviorally relevant visual input under natural conditions. While parametric stimuli, such as the chirps and moving bars, provide a powerful and interpretable framework for classifying retinal inputs, they only sparsely sample the high-dimensional visual space encountered in natural environments. Conversely, natural movies capture the statistics of ethologically relevant vision but are difficult to analyze exhaustively, motivating the need for a model that can learn from naturalistic input while remaining interpretable in terms of established functional dimensions.

To address this, we built a functional “digital twin” of retinal input to the SC: a deep dynamic model that learns the stimulus–response transformation of RGC axonal boutons directly from natural movies, while generalizing to parametric and novel visual stimuli not used during training (Fig. 4a). Conceptually, this approach creates an *in silico* replica of retinal drive to the SC, enabling large-scale *in-silico* experiments that would be impractical—or impossible===*in vivo*.

**Fig. 4.**
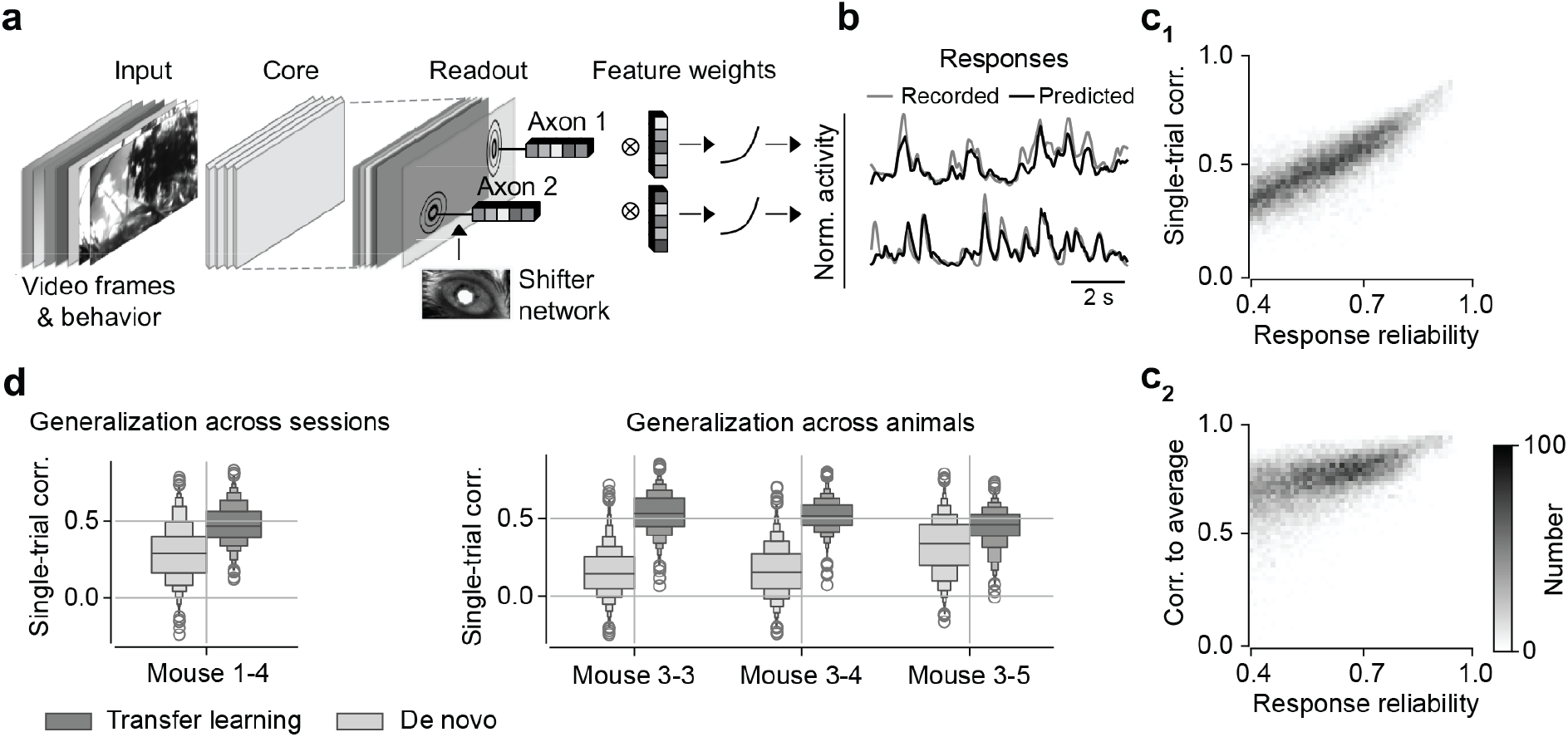
A deep dynamic model predicts responses of mouse retinal axons. **a**, Schematic of the predictive model. Natural movie frames and behavioral variables were provided as inputs to a three-dimensional convolutional core that extracted shared spatiotemporal features. A scan-wise Gaussian readout mapped core features to individual axonal boutons, while a shifter network dynamically adjusted readout positions to compensate for eye movements during stimulus presentation. **b**, Example responses of two axonal boutons to natural movie stimuli (gray, recorded activity; black, model prediction), shown as normalized activity traces. **c**, Two-dimensional histograms summarizing model performance across 17,272 functional units from ten experimental scans. Only units meeting minimum quality thresholds were included for model training: reliability (mean leave-one-out correlation across trial repetitions, collapsing across trial order) *>* 0.4, and stability (absolute slope of leave-one-out correlations across successive trials) *<* 0.2. Top: single-trial correlation between predicted and recorded responses versus response reliability. Bottom: correlation between predicted responses and trial-averaged responses versus response reliability. **d**, Box plots comparing single-trial correlation for transfer learning (dark gray) and *de novo* (light gray) models, when testing generalization across recording sessions within the same animal (left) and across animals (right). Transfer learning consistently outperformed *de novo* training in both scenarios.

The model architecture consisted of a three-dimensional convolutional core shared across boutons, with factorized spatial and temporal kernels extracting shared spatiotemporal features from visual input. Individual bouton responses were read out via bouton-specific Gaussian read-outs, with shifter networks compensating for eye move-ments during stimulus presentation (Sinz et al., 2018; Lurz et al., 2020a). The convolutional core captured common retinal computations, while the readouts encoded bouton-specific functional fingerprints.

We trained the model on responses to natural movies, reserving repeated movie presentations exclusively for evaluation. Training on 10 experimental recordings from three animals yielded high predictive performance on held-out data, with a median single-trial correlation of 0.500 and a median correlation to trial-averaged responses of 0.761 (Fig. 4b,c; Supple. Fig. 6), indicating that the model accurately captured both trial-to-trial variability and stimulus-locked response structure.

A critical requirement for a functional digital twin is generalization beyond the specific data used for training (*transfer learning)*. To test this, we first assessed transfer across recording sessions within the same animal. A model trained on three recordings from one animal and fine-tuned on a fourth recording from the same animal—while keeping the convolutional core fixed—outperformed a *de novo* model trained solely on the fourth recording (Fig. 4d; *p <* 0.0001 for a paired two-sided permutation test with 10,000 repeats). This result indicates that the learned core captures stable, recording-invariant retinal computations. We next evaluated generalization across animals by training a shared convolutional core on data from two animals and fine-tuning readouts on a third. Again, transfer learning consistently yielded higher single-trial correlations than *de novo* training across all scans (Fig. 4d; *p <* 0.0001 for a paired two-sided permutation test with 10,000 repeats), demonstrating that the model captures retinal computations that generalize across individuals.

Together, these results establish a high-performance functional digital twin of retinal input to the SC. In the following sections, we show that this model generalizes to classical parametric stimuli and enables predictive analyses of retinal encoding under behaviorally relevant visual conditions.

### The model captures anatomical structure and generalizes across stimulus domains

Having established the model as a functional digital twin, we next examined whether its learned internal representations captured biologically meaningful structure and generalized across stimulus domains. To probe the feature selectivity encoded by the model, we computed dynamic most exciting inputs (MEIs) for the recorded boutons (Walker et al., 2019; Bashivan et al., 2019; Ponce et al., 2019; Höfling et al., 2024). This approach identifies the nonlinear spatiotemporal receptive fields of individual boutons. Singular value decomposition of these MEIs revealed temporal components with complex, multiphasic profiles containing multiple positive and negative peaks, while the spatial components often extended beyond simple center–surround or small, localized receptive field structures (Fig. 5a). Many MEIs exhibited spatial–temporal patterns suggestive of complex motion selectivity, consistent with prior work (Höfling et al., 2024; Li et al., 2025; Jo et al., 2023).

**Fig. 5.**
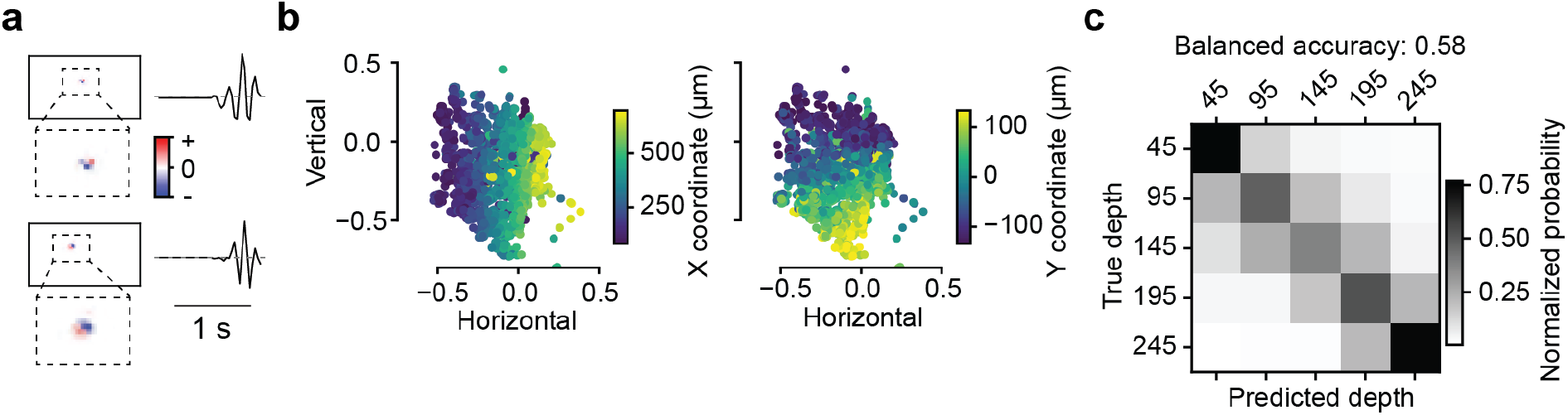
The model captures anatomically meaningful structure. **a**, Dynamic most exciting inputs (MEIs) computed for two example axonal boutons. Insets show zoomed-in spatial components of the MEIs, while traces depict the corresponding temporal components, illustrating the spatiotemporal stimulus features that maximally drive individual model units. **b**, Learned spatial positions of Gaussian readouts for all boutons from one representative scan, projected into model coordinate space and color-coded by experimentally measured cortical coordinates along the vertical (left) and horizontal (right) axes. The smooth gradients indicate that the model’s learned readout positions align with the retinotopic organization of the recorded data. **c**, Confusion matrix comparing true versus predicted cortical depth of axonal boutons, based on a classifier trained on readout feature weights (cf. Fig. 4a). Values are normalized such that each row sums to 1.0. The diagonal structure reflects above-chance prediction of bouton depth from functional representations alone (balanced accuracy = 0.58).

We next asked whether the model’s internal parameters reflected the anatomical layout of recorded RGC axon terminals within the recorded brain volume. We focused on the Gaussian readouts, which are parameterized by a spatial position and a set of feature weights. Visualization of the learned receptive field centers revealed clear gradients along the horizontal and vertical axes of the stimulus, indicating strong alignment with experimentally recorded cortical coordinates (Fig. 5b).

We then tested whether the learned readout weights— forming a 64-dimensional functional fingerprint—encoded additional anatomical information beyond retinotopy. A logistic regression model trained to predict bouton cortical depth from these weights achieved a balanced accuracy (average of recall obtained on each class) of 0.58, substantially exceeding the chance level of 0.2 (Fig. 5c), demonstrating that the model captured laminar organization from functional data alone.

Beyond the anatomical layout, we examined whether the model—trained exclusively on natural movies— generalized to classical parametric stimuli. The model accurately predicted responses to both full-field chirp and moving bar stimuli across all three mice (Fig. 6a,b; Supple. Fig. 7a,b). Preferred directions and orientations derived from model predictions closely matched those obtained from experimentally recorded responses (Fig. 6c). To further assess the biological relevance of the learned representations, we related readout feature weights to specific response properties. A ridge regression model trained on these weights predicted the On-Off index (OOi) with an *R*^2^ of 0.56 (Fig. 6d), and OOi values varied smoothly when boutons were projected onto the first two principal components of the readout weights (Fig. 6e). Finally, a logistic classifier based solely on readout weights recovered bouton response types identified by Gaussian mixture model clustering with a balanced accuracy of 0.73, far exceeding the chance level of 0.05 (Fig. 6f). Together, these results show that the digital twin learns interpretable, anatomically grounded representations that generalize from naturalistic visual input to classical parametric descriptions of retinal function.

**Fig. 6.**
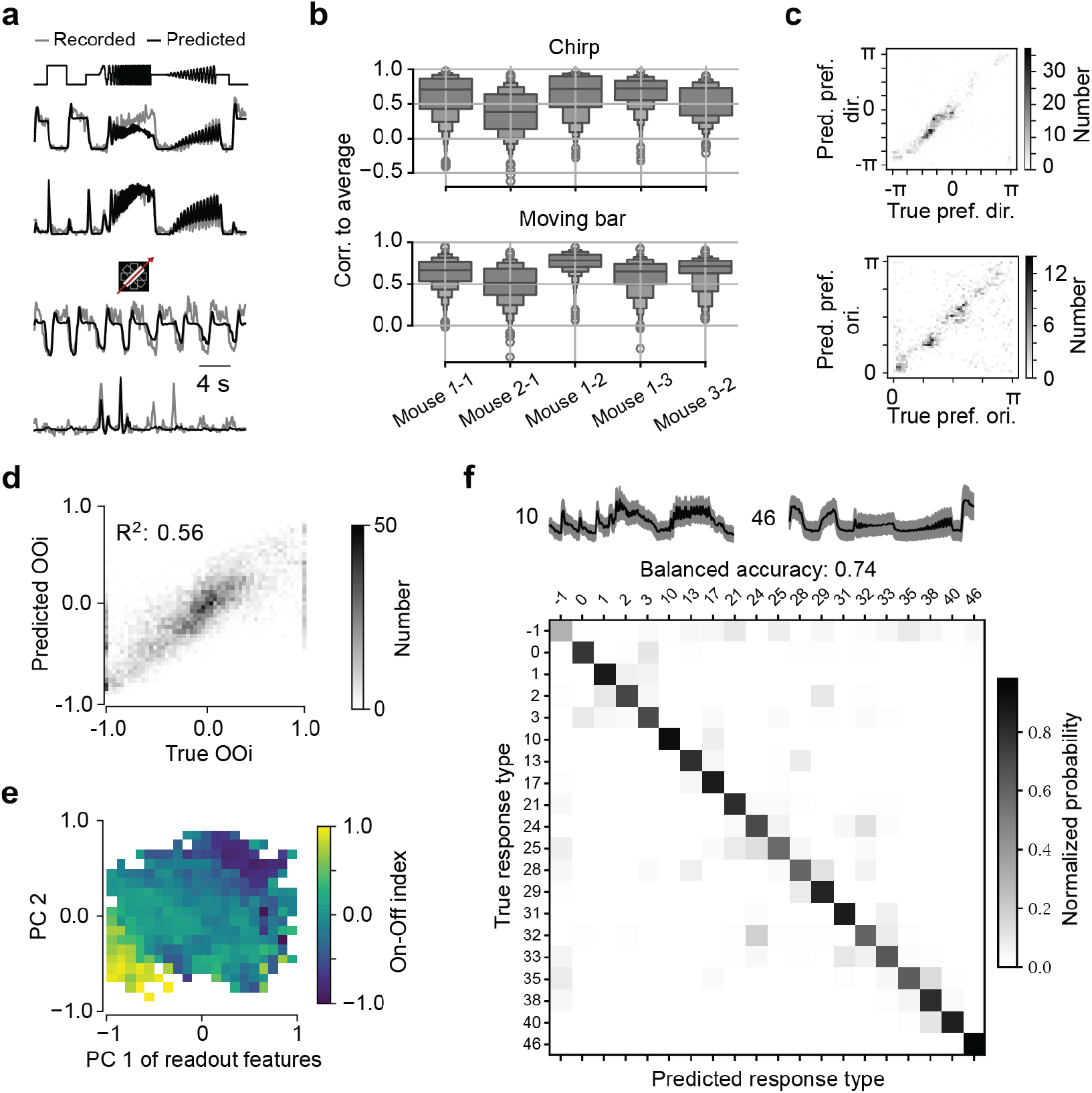
Prediction of functional properties computed from responses to ‘chirp’ and moving bar. **a**, Example neuronal responses to full-field ‘chirp’ (upper) and moving bar (lower), comparing recorded traces (gray) with model predictions (black). **b**, Model performance quantified as correlation to the trial-averaged response for the full-field ‘chirp’ (upper) and moving bar (lower). Values close to the noise ceiling indicate the model captures the majority of explainable variance. **c**, Upper: 2D histogram of predicted vs. true preferred direction for boutons in one scan field; lower: corresponding histogram for preferred orientation. Concentration along the diagonal reflects accurate tuning predictions. **d**, 2D histogram of predicted vs. recorded On-Off index (OOi) across the bouton population. **e**, 2D heatmap of boutons positioned by the first two PCs of their readout feature weights, colorized by OOi. Clustering in feature space suggests the model captures functionally meaningful structure. **f**, Upper: mean ‘chirp’ responses (black, mean; shaded gray, standard deviation) for two exemplary response types identified from readout feature weights; lower: confusion matrix comparing predicted response types against GMM clustering on measured responses. Each row is normalized to sum to 1.0.

**Fig. 7.**
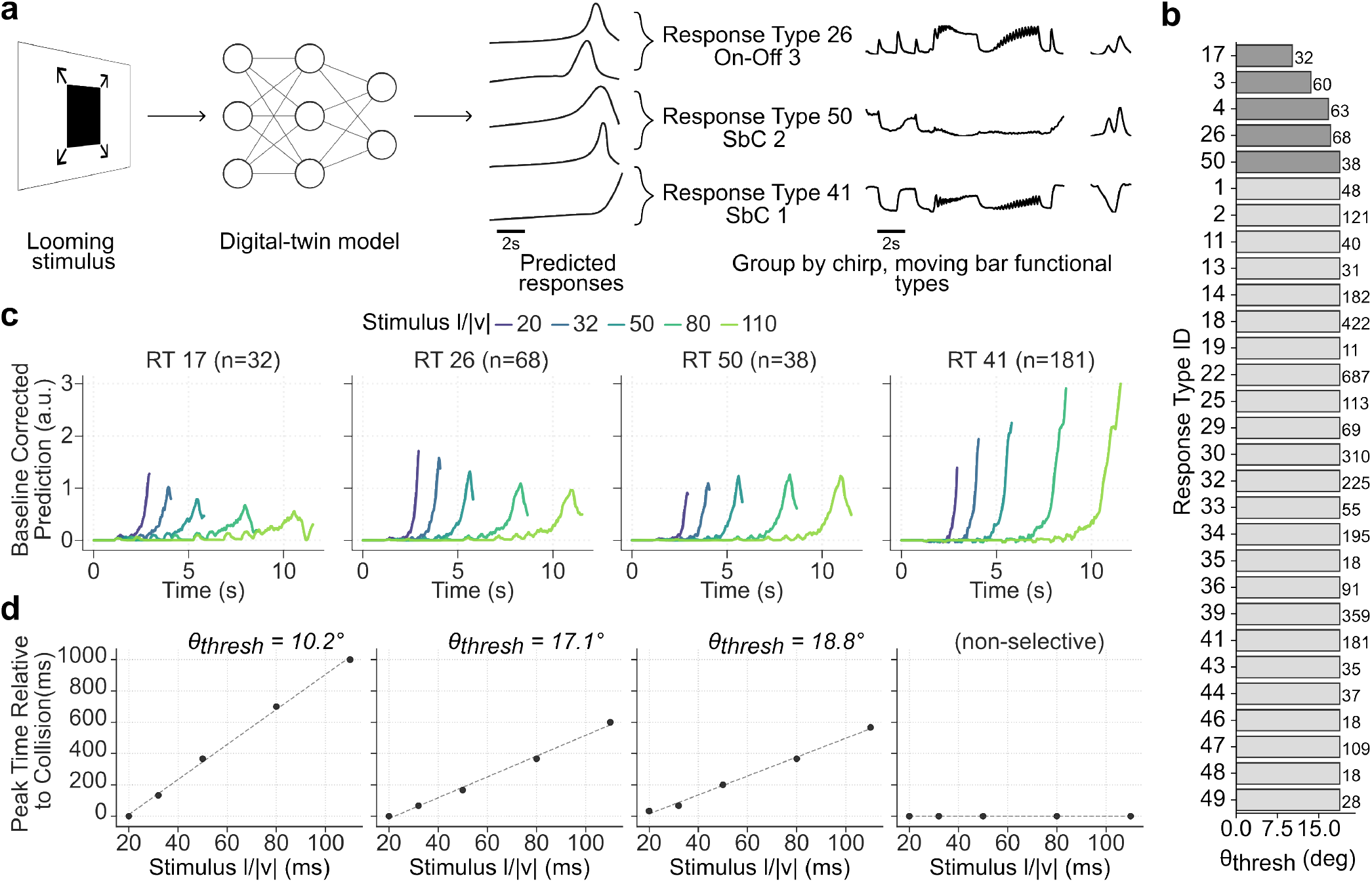
Using the digital twin to predict encoding of looming stimuli. **(a)** Schematic: looming stimuli were presented *in silico* to the trained “digital twin” model (presented in Fig. 4a) to generate predicted bouton responses, which were then aggregated by functional groups defined from ‘chirp’ and moving bar responses. **b**, Angular threshold *θ*_thresh_ of functional groups to looming stimuli. Number of neurons indicated to the right of bars. Looming selective groups with *R*^2^ *>* 0.95 are shown in dark gray; all other non-selective groups are shown in light gray. Groups with less than 10 neurons were excluded. **c**, Predicted mean responses of selected functional groups to looming stimuli with different *l/*|*v*| values (colored traces; legend shown above the plots). Response Type (RT) indices correspond to Fig. 2; *n* denotes number of boutons per group. **d**, Relation between stimulus *l/*|*v*| and peak time relative to collision. Dots indicate the time of peak response relative to collision, *t*_peak_, for each *l/*|*v*|, and dashed lines show linear fits of *t*_peak_ versus *l/*|*v*| for looming-responsive groups. The inferred angular threshold *θ*_thresh_ is reported for groups exhibiting consistent scaling (first three panels), whereas an example non-selective response group shows no systematic relationship.

### Predicting responses to behaviorally relevant looming stimuli

A key advantage of the functional digital twin is that it can be driven *in silico* with arbitrary visual input, enabling systematic tests of stimulus encoding that would be difficult to perform experimentally at scale. We focused on looming stimuli because they are an ethologically-relevant signal driving rapid escape or freezing behaviors, and have been studied extensively in insects and vertebrates including mice (e.g. Yilmaz and Meister, 2013). Inspired by work characterizing looming-sensitive neurons in insects (Gabbiani et al., 1999), we tested whether responses follow a canonical signature of collision detection: peak response time scaling linearly with the stimulus parameter *l/*|*v*| (object half-size divided by approach speed), consistent with detection at a fixed angular threshold on the retina. This relationship provides a compact signature of collision-sensitive computations: across different *l/*|*v*| values, the peak response time shifts in proportion to *l/*|*v*|, consistent with detection at a fixed angular size.

We used the trained digital twin to predict bouton responses to looming stimuli spanning multiple *l/*|*v*| values (Fig. 7a). Predicted responses were then grouped according to functional response types defined by chirp and moving bar stimuli, allowing us to compare looming sensitivity across retinal response types. To quantify looming selectivity at the level of response types, we followed the classic parametric analysis used in the locust literature and examined whether the time of peak response relative to collision, *t*_peak_, varied linearly with *l/*|*v*| within each cluster. We identified a small subset of looming-selective response types—comprising Off, On, and Suppressed-by-Contrast types—in which *t*_peak_ scaled approximately linearly with *l/*|*v*|, yielding angular thresholds ranging from approximately 10° to 15° (Fig. 7c,d; Methods). The remaining response types showed no clear linear relationship between *t*_peak_ and *l/*|*v*|, resulting in substantially larger and less consistent angular threshold estimates (Fig. 7b). Together, these results illustrate how the digital twin can be used as an *in silico* assay to narrow down candidate retinal input channels for collision detection, identifying a discrete subset of functionally defined response types with physiologically plausible looming selectivity from among the full diversity of retinal inputs to the SC.

## Discussion

Our study reveals that the mouse SC receives input representing nearly the complete functional repertoire of RGC types characterized in *ex vivo* retinal recordings, preserving a high-dimensional feature space that changes with SC anatomical depth. By capturing these dynamics in a CNN trained exclusively on natural movie responses, we created a functional digital twin that successfully generalizes to predict responses to unseen parametric stimuli, including well-established frequency and contrast chirps and moving bars. This cross-stimulus predictive power validates the model as a reliable *in silico* exploration tool. We demonstrate this capability by replicating looming selectivity—an ecologically relevant, out-of-training-set signal—in the model.

### Preservation of retinal feature diversity in the SC

Anatomically, the mouse SC receives input from the vast majority of RGCs, with approximately 90% of RGCs projecting to the SC (Ellis et al., 2016). Further, functionally distinct RGC populations stratify at stereotyped depths within the superficial SC (Huberman et al., 2008; Hong et al., 2011), suggesting that the SC receives a structurally ordered map of parallel retinal channels. However, while these studies established broad access to retinal signals, they did not determine whether the full functional diversity of retinal output is preserved at the level of SC input, or how this diversity is structured across depth within the collicular circuit.

Previous functional studies have provided partial answers, but often focused on specific retinal subtypes. Targeted anatomical and physiological experiments mapped the projection patterns and response properties of individual or selected RGC types across SC layers, revealing clear relationships between cell type, laminar position, and tuning properties (Reinhard et al., 2019; Kühn et al., 2025). While these studies provided important mechanistic insights, their focus on specific RGC classes precluded an unbiased assessment of the full diversity of retinal signals entering the SC.

More recently, large-scale recordings have provided increasingly comprehensive views of visual processing in the SC. Using Neuropixels recordings from retinal axons in the SC, Sibille et al. (2022) showed that multiple, functionally distinct retinal pathways provide precise and reliable input to collicular neurons, but that the diversity of functionally separable inputs was lower than expected from *ex vivo* classifications of RGC types (Baden et al., 2016; Goetz et al., 2022). Complementing this approach, **?** imaged RGC axonal boutons in the superficial SC and found that retinal input signals are modulated by arousal state. Borges et al. (2025) imaged RGC axon boutons in the superficial SC and showed that retinal inputs encode saliency and exhibit strong, context-dependent tuning, consistent with retinal center–surround mechanisms; however, this work did not examine the diversity or completeness of retinal functional types contributing to SC input. In parallel, Li and Meister (2023b) performed large-scale calcium imaging of postsynaptic SC neurons and identified 24 functional SC cell types—substantially fewer than the >45 RGC types described in the retina (Baden et al., 2016; Goetz et al., 2022). They further showed that individual SC neurons typically pool inputs from multiple RGC types, indicating substantial convergence and feature recombination within collicular circuits (also see Lee et al., 2020). Together, these studies indicate that SC neuronal representations may potentially be less complex and lower dimensional than retinal output, but leave open whether this reduction reflects a consequence of sampling biases, reduced retinal input or instead emerges through circuit-level transformations applied to a richly structured retinal input.

Here, we addressed this question by labeling essentially all RGCs in the retina and selectively recording their axonal activity in the SC. Using two-photon imaging, we observed a strikingly rich diversity of retinal input responses across SC layers. By aligning *in vivo* bouton responses with large-scale *ex vivo* retinal datasets in a shared latent space, we found that the SC receives a near-complete representation of retinal functional diversity. Compared to previous axonal recordings using extracellular electrophysiology (Sibille et al., 2022), this difference likely reflects the ability of two-photon imaging to densely sample small-diameter axons and synaptic boutons that may be underrepresented in extracellular recordings.

Our analysis identified approximately 50 functional response types that were systematically organized across SC depth. While the exact number of types depends on methodological choices, the strong correspondence with *ex vivo* retinal types supports the conclusion that parallel retinal feature channels are largely preserved at the level of SC input. In addition, we observed a subset of response types that were preferentially detected in the SC and are not typically defined in *ex vivo* retinal recordings. Rather than reflecting noise or misclassifications, these clusters may arise from physiological processes present only in the intact *in vivo* preparation, such as local presynaptic mechanisms (Mooney et al., 1994; Govindaiah et al., 2020) that dynamically shape retinal axon terminal activity in the SC, or state-dependent neuromodulation (Schröder et al., 2020; Savier et al., 2019) that may differentially affect retinal processing under awake behaving conditions.

Taken together, our results demonstrate that the SC receives a near-complete and richly structured representation of retinal functional diversity at the level of its input. Rather than indicating early selection of retinal signals, our findings suggest that the reduction in diversity observed at the level of SC neuronal responses (Li and Meister, 2023a) emerges downstream, through convergence and recombination of retinal inputs by local collicular circuits. This organization provides collicular circuits with access to a broad repertoire of retinal feature channels, forming a flexible substrate upon which subsequent circuit-level transformations can operate.

### Bridging functional cell type classification with ecological relevance

A persistent challenge in neuroscience is reconciling functional cell-type classifications derived from standardized, parametric stimuli with representations shaped by the statistics of natural input. Large-scale atlases based on artificial probes such as frequency chirps, moving bars, and flashed spots have provided a rigorous and reproducible framework for defining RGC diversity (Baden et al., 2016; Goetz et al., 2022). However, because these stimuli are only weakly constrained by natural scene statistics, it has remained unclear how the resulting functional dimensions relate to ecologically grounded visual representations.

To bridge this gap, we utilized a deep learning model of retinal input to the SC as an *in silico* testbed. We interrogated this model—which was optimized solely to encode natural movie responses—using standard parametric stimuli to determine if the neural computations required for natural vision spontaneously give rise to canonical tuning properties.

A key result is that this model generalized robustly to classical parametric probes. Despite never being trained on artificial stimuli, the model reproduces well-established response features measured with chirps and moving bars— features that form the basis of widely used retinal and SC functional taxonomies (e.g. Baden et al., 2016; Lee et al., 2020). This finding parallels recent work by Höfling et al. (2024), who showed that models trained on natural movie responses can accurately capture retinal computations traditionally characterized using artificial stimuli. That response patterns to parametric stimuli emerge from a model trained exclusively on natural movies supports the notion that such tuning properties capture computations that extend beyond the artificial stimulation paradigm.

This perspective is consistent with a growing body of work in visual cortex demonstrating that models trained on natural stimuli can reproduce—and thereby reconcile— classical results obtained with artificial stimuli. In particular, recent modeling studies have shown that representations learned from natural scenes capture response properties previously identified using highly controlled stimulus paradigms, providing a unified account of sensory processing (Fu et al., 2024). By extending this insight to the retinal output, our analyses suggest that the alignment between natural statistics and parametric tuning is a fundamental organizing principle established at the earliest stages of visual processing.

Importantly, the resulting model is intended as a shared community resource. Beyond serving as a predictive tool, this framework acts as a high-throughput *in silico* exploration tool. In the spirit of open modeling efforts such as OpenRetina (D’Agostino et al., 2025), we make our trained model publicly available to enable further studies of the functional properties of retinal input signals to the SC. Because the model generalizes across stimulus classes, it enables the systematic dissection of non-linear integration properties and the screening of vast stimulus spaces that are often intractable in biological experiments. We anticipate that this model will support future experimental and theoretical work aimed at linking established functional descriptions to ecologically grounded representations of visual input.

### Limitations and challenges of the approach

While our framework provides a principled method to reconcile functional diversity across *ex vivo* retinal recordings and *in vivo* SC inputs, several limitations warrant consideration.

Our conclusions rely on the representativeness of the sampled RGC population. Sampling biases cannot be fully excluded, particularly if specific RGC types expressed the calcium indicator weakly or were less amenable to optical detection. However, histological co-staining for GCaMP and a pan-RGC marker confirmed that the vast majority of RGCs were labeled, making systematic omission of specific types unlikely. Subtle variations in labeling efficiency may nonetheless influence relative cluster occupancy and should be considered when interpreting quantitative type distributions.

The choice of 50 clusters does not imply a ground-truth estimate of the number of distinct RGC types projecting to the SC, but rather reflects a balance between resolving meaningful functional variability and avoiding spurious structure arising from technical sources such as session-to-session variability. Our objective was not to establish a one-to-one correspondence between *in vivo* inputs and *ex vivo* types, but to verify whether the major functional motifs governing retinal output are preserved in SC input.

Third, aligning these datasets entails fundamental challenges due to distinct experimental regimes. Despite efforts to match stimuli, the recordings differed in preparation (*in vivo* vs. *ex vivo*), optical path, and stimulus size and geometry. Consequently, our contrastive alignment constructs a shared functional coordinate system rather than asserting exact equivalence. By focusing on distances and overlaps within this latent space, we identify corresponding functional groups without forcing direct cell-to-cell matches. This approach necessarily involves a trade-off: while the alignment effectively suppresses technical variability, it risks discarding domain-specific biological signals alongside the noise. While most response features were faithfully reconstructed from the latent space, a small subset of outliers suggests that the embedding is not uniformly precise for all response profiles.

Finally, our analysis of looming responses serves as a proof-of-concept for the model’s predictive capacity. These results should be viewed as hypothesis generation rather than ground truth; we have not yet experimentally validated the specific dynamics of the predicted looming responses. However, the predicted relationship between response peak timing and time-to-collision is qualitatively consistent with classic studies of collision-detecting circuits in locusts (Gabbiani et al., 1999). This convergence suggests the model captures meaningful biological structure, while underscoring the need for targeted experiments to verify that these specific computations are implemented in the retino-collicular pathway.

### From retinal diversity to collicular function

The observation that the mouse SC receives input from nearly the full complement of RGC types identified in *ex vivo* retinal recordings raises a critical question: what is the computational advantage of maintaining such high-dimensional input in a subcortical structure? Our findings suggest that the rodent SC operates not merely as a relay for reflex generation, but as a sophisticated interface for parallel feature processing (Lee et al., 2020; Li and Meister, 2023b; Wang et al., 2010; de Malmazet et al., 2024; DePiero et al., 2024). This distinction is particularly relevant when viewing the colliculus through a comparative lens. In primates, the geniculostriate pathway (Retina→ LGN→ V1) is the dominant conduit for high-resolution feature analysis, while the SC is often conceptualized primarily as a center for saccadic control and spatial attention (Sparks, 1986; Krauzlis et al., 2013). In contrast, our results highlight that the rodent SC accesses a feature space of comparable richness to that of the cortex, implying that complex visual feature extraction can be performed by the midbrain directly—allowing the SC to orchestrate behavior without strictly requiring cortical top-down feature refinement. Notably, recent evidence suggests that even in primates, the SC contributes to visual processing beyond its classical roles, including object detection and abstract categorization (Hafed et al., 2023; Peysakhovich et al., 2024), hinting that its capacity for feature-level computation may be a conserved property across mammalian evolution.

Finally, the retinal axon model established here provides a critical foundation for future computational dissection of the mouse collicular circuit. Cascading this input characterization with models of downstream SC circuitry— incorporating lateral inhibition or projection-specific dendritic integration—opens a principled path toward end-to-end *in silico* analysis of how the retina’s feature dictionary is transformed into behavioral output, bridging the gap between sensory encoding and motor control.

## Methods

### Surgery and viral delivery

All experimental procedures complied with NIH guidelines and were approved by the Baylor College of Medicine Institutional Animal Care and Use Committee (protocol AN-8132). AAVs expressing GCaMP8m under the hSyn promoter were delivered via intraocular injection into anesthetized mice, resulting in labeling of retinal ganglion cells (RGCs) and their axons projecting to the superior colliculus (SC). To implant the cranial window, the right arm of the transverse sinus was ligated and sutured to allow access to the SC. A custom 3D-printed cranial window (3 mm diameter) was subsequently implanted over the SC, with overlying cortical tissue carefully displaced laterally but not aspirated, preserving tissue integrity while allowing stable optical access. Post hoc histological analysis confirmed accurate window placement and the absence of a detectable immune response, as assessed by Iba1 staining (Fig. 1b Holtmaat et al., 2009). The preparation supported repeated imaging sessions for up to six months.

### Two-photon calcium imaging and data collection

Calcium signals from RGC axonal boutons were recorded using two-photon imaging in awake, head-fixed mice (n = 3, aged 6–13 months at the time of recording). A total of nine recordings were performed across up to five optical planes spanning 20–300 µm depth from the SC surface, with field sizes of 270×400-619 µm (field areas of 0.108 - 167 mm^2^) at a recording resolution of 0.54 - 0.68 µm/px. Each scan covered 0.54 - 0.836 mm^2^, yielding between 2,275 and 8,029 segmented masks per scan (mean: 5,471). Functional recordings were motion corrected over time; fluorescence signals (rather than deconvolved traces) were used for all subsequent analyses. Regions of interest (ROIs) corresponding to individual axonal boutons were automatically segmented from the motion-corrected fluorescence movies using constrained non-negative matrix factorization within the CaImAn pipeline, and were subsequently inspected to exclude artifacts. Eye position and pupil dynamics were monitored continuously via an infrared camera throughout each session. In total, the dataset comprised recordings from 229,774 individual RGC axonal boutons sampled at 8 Hz from n = 42 recordings. All animals were drug- and procedure-naïve prior to these experiments. A detailed description of the imaging setup, laser parameters, and signal extraction pipeline is provided in Franke et al. (2022) and Wang et al. (2025).

### Visual stimulation

Two parametric stimulus classes were presented (Baden et al., 2016): (i) a full-field chirp consisting of a bright step followed by sinusoidal modulations of increasing temporal frequency and contrast (15 trials); and (ii) a bright bar (0.3×1 mm) moving at 1 mm s_−1_ in eight directions on a dark background (10 trials). In separate sessions, natural movie clips were shown interleaved with parametric stimuli (for details, see Wang et al., 2025; Turishcheva et al., 2023). Visual stimuli were displayed on an LCD monitor (60 Hz refresh rate, 31.1 cm x 55.3 cm monitor placed 15 cm from their left eye) calibrated for luminance using a photodiode and luminance meter prior to each session.

### Pre-processing and quality control

Functional responses and behavioral signals (pupil size, locomotion) were resampled to 30 Hz using linear spline interpolation. Response stability was assessed by computing leave-one-out correlations across repeated stimulus presentations and quantifying monotonic drift across the session; boutons exhibiting drift above the threshold (see below) were excluded. Response reliability was quantified as the mean leave-one-out correlation across trials. For clustering of parametric stimulus responses, we used axons with a response reliability greater than 0.3 for the chirp stimulus and greater than 0.4 for moving bars, yielding 66,029 out of 201,352 boutons (n = 37 recordings, n = 3 mice). For model training, we selected boutons with an oracle correlation to repeated test movies greater than 0.4 and a drift slope across repeats smaller than 0.2, resulting in 17,272 out of 63,747 boutons (n = 10 recordings, n = 3 mice). Throughout, we use the terms “bouton” and “axon” interchangeably to refer to a single region of interest defined by functional similarity of pixel responses; this does not necessarily correspond to a single anatomical axonal bouton.

### Data pre-processing for clustering

For the parametric stimuli, we applied quality filtering to the recorded responses based on their scan quality score, using thresholds of 0.3 and 0.4 for the chirp and moving bar stimuli (Fig. 1a), respectively. Artifact responses were identified as described previously (Schmors et al., 2025) and removed before clustering. We also excluded a small number of responses with sharp negative deflections that are incompatible with the slow off-kinetics of GCaMP (Zhang et al., 2023) (Fig. 1b, bottom response). They display inverted fast-Off, slow-On kinetics, which previous studies have indicated are not reported by GCaMP sensors (Wei et al., 2020). Thus, we interpret these sharp negative deflections as artifacts that are a consequence of the background subtraction performed by CaImAn segmentation (Giovannucci et al., 2019) being applied to low-signal ROIs. This resulted in a dataset of 66,029 individual axon boutons. For each bouton, we trim the first second, z-score single trials, average across trials with baseline correction (first 200 ms to zero), and normalize to maximum absolute value of 1.

### Clustering

We clustered the functional responses to the parametric stimuli (chirp and moving bar) using a Mixture of Gaussians model (GMM). For feature extraction, we applied sparse principal component analysis (sPCA) with sparsity controlling parameters of *α* = 20 and 40 for the chirp stimulus and moving bar responses, respectively. For the moving bar responses, we concatenated responses to the eight directions instead of extracting only the mean time course across directions as done in Baden et al. (2016). This allows for automatic identification of both direction-selective and orientation-selective response profiles. The resulting temporal features were then used for GMM clustering with a diagonal covariance matrix, which initially yielded n=82 clusters based on the minimum Bayesian Information Criterion (BIC) Baden et al. (2016). To ensure cluster quality, we conditioned clusters to have an intra-cluster correlation *r >* 0.7 (leave-one-out correlation) in either bar or chirp responses. We further excluded any cluster in which more than 70% of ROIs originated from the same depth or the same scan field, ensuring that retained clusters reflect functional response profiles present across multiple scans rather than scan- or depth-specific batch artifacts. This filtering resulted in n=61 clean clusters, which were then merged into *n* = 50 final clusters for visualization purposes. We arranged clusters in a tree according to their similarities using the correlation distance (1 − *r*_Pearson_) and optimal leaf ordering by minimizing the distances between adjacent clusters in the tree.

### Characterization of responses to parametric stimuli. Direction and orientation selectivity

To quantify the direction and orientation selectivity of the Ca^2+^ responses, we extracted temporal and directional components of the moving bar responses according to Baden et al. (2016). In short, we averaged the responses across trials to obtain a mean response matrix **M** of size *T*×*D* (time bins by directions, *T* = 32, *D* = 8). We performed singular value decomposition (SVD) on **M** to decompose the responses into temporal and directional components: **M** = **USV**^*T*^, with the first column of **U** representing the temporal component, and the first row of **V** representing the directional tuning. Preferred direction and preferred orientation were computed as the phase angles of complex vector sums obtained by projecting the normalized directional tuning curve **v** onto single-frequency and double-frequency complex exponentials, respectively (for implementation details see Baden et al. (2016)). The direction selectivity index (DSi) and orientation selectivity index (OSi) were quantified as the magnitude of the respective vector sums. Statistical significance of direction and orientation selectivity was assessed using a permutation test with 100 shuffles, where trial labels were randomly shuffled to generate a null distribution.

### Response transient index

The response transience index (RTi ∈ [0, 1]) quantifies the response kinetic of a visual cell during the first step response of the chirp stimulus. Response with sustained characteristics have a RTi = 0, whereas transient responses with a decay back to baseline have RTi = 1. The RTi was computed as

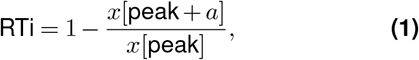

where *peak* defined the time point of peak response and *a* = 400 ms the response following the peak.

### On-Off index

Preference to On and Off light stimulation was measured as

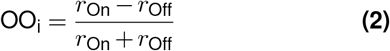

where *r*_*On*_ and *r*_*Off*_ are defined as the responses to the step component of the chirp stimulus (the first 200 ms of the On step and the first 200 ms of the Off step).

### Suppression index

For each cell, the suppression index (SI) was computed by comparing the absolute negative area under the curve (AUC_neg_) to the total response area (AUC_neg_+AUC_pos_) of the ‘chirp’ and moving bar responses. Response traces were clipped to isolate negative and positive components prior to numerical integration. The SI was defined as:

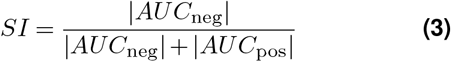

### Variational autoencoder with domain-adversarial learning: Model architecture

To learn a shared representation across *ex vivo* RGC and *in vivo* bouton responses, we used a variational autoencoder (VAE) combined with domain-adversarial learning. The model was designed to preserve stimulus-driven response structure while minimizing dataset-specific domain information.

The encoder consisted of two fully connected layers (hidden layer size 128) with ReLU nonlinearities and dropout. From the encoder output, separate linear layers predicted the latent mean *µ* and log-variance log*σ*^2^. Latent samples were obtained via the reparameterization trick. The decoder mirrored the encoder, mapping latent variables back to the original response space using fully connected layers and ReLU activations, and was optimized using mean-squared reconstruction error.

To encourage domain invariance, a domain discriminator operated on the latent representation. The discriminator was a multilayer perceptron with two hidden layers (size 128, ReLU activations) and a binary output predicting dataset identity (RGC vs. Bouton). A gradient reversal layer was placed between encoder and discriminator so that the encoder learned to reduce domain-predictive information in the latent space.

To preserve functional information, we included dataset-specific classification heads predicting RGC group or bouton response type identity from the latent representation using cross-entropy loss.

### Variational autoencoder with domain-adversarial learning: Model training and evaluation

Models were trained using the Adam optimizer with weight decay. The loss function was a weighted sum of reconstruction loss (mean-squared error), Kullback–Leibler divergence, type-classification loss, and domain-adversarial loss. The strength of gradient reversal was gradually increased during training following a sigmoidal schedule.

To avoid domain imbalance, each mini-batch contained equal numbers of RGC and bouton samples. Training proceeded for a fixed number of epochs, and analyses used the mean of the posterior latent distribution. To quantify domain alignment, we trained a logistic-regression domain classifier on latent representations using stratified cross-validation. Performance was reported as classification accuracy. Reduced domain-classification performance indicated successful domain invariance. Reconstruction quality was assessed using the mean-squared error between original and reconstructed responses, computed per cell and averaged within cell types.

### Variational autoencoder with domain-adversarial learning: Data pre-processing

RGC and bouton response traces were first resampled to a common temporal resolution using linear interpolation. All traces were interpolated to 260 time points over a shared stimulus window to ensure temporal alignment across datasets. To standardize response amplitudes while preserving temporal structure, each response trace was normalized by its maximum absolute value. For a response vector **r**_*i*_, normalization was performed as:

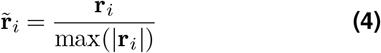

such that all responses lay within the range [*−*1, 1].

### Latent-space mapping between RGC groups and bouton response types

To quantify functional correspondence between RGC groups and bouton response types, we computed mean latent representations (‘type centroids’) for each group or response type. Similarity between RGC and bouton centroids was measured using cosine similarity, yielding a cross-dataset similarity matrix.

For each RGC group, the bouton response type with the highest cosine similarity was taken as the best functional match. This approach allowed us to identify structured correspondences between dataset response types in the shared latent space.

### Model architectures

We used a dynamic network consisting of a 3D factorized convolutional core (Höfling et al., 2024), Gaussian readouts (Lurz et al., 2020b), and shifter modules (Sinz et al., 2018) to model stimulus-response functions.

The core included three sequential structures with a spatial convolutional layer (64 output channels) and a temporal convolutional layer (64 output channels) followed by a batch normalization layer and an exponential linear unit (ELU) function. The first spatial kernel was 1×11×11 and the other two were 1×5×5. Similarly, the first temporal kernel was 11×1×1 and the other two were 5×1×1.

To estimate neuronal activity for each axonal bouton from the core output tensor (*w*×*h*×*c*, width x height x channels), we performed generalized linear regression with a learnable weight tensor (*c*×*w*×*h*) and an ELU function offset by one (ELU + 1) to enforce positive response. To reduce the learnable parameters in this regression problem, we adopted Gaussian readout computing a 2D Gaussian distribution *N*(*µ*, Σ) (*µ*, mean location of width and height in the core output tensor; Σ, uncertainty of the location) for each bouton’s receptive field position (Lurz et al., 2020b). Parallel to the position learning, the model learned the feature weights of size *c* per bouton. For the model with access to the recorded 2D cortex coordinates of each bouton, we used a multilayer perceptron (MLP) with a size of 2 30 2, shared by all boutons in one session, to map the coordinates to the mean location *µ* (Bashiri et al., 2021). To account for the mouse eye movements during stimulus presentation, we employed a shifter network using the pupil center to shift the model neuron’s receptive field center *µ* (Sinz et al., 2018). The shifter is an MLP consisting of three fully connected layers with *n* = 5 hidden features and a *tanh* nonlinearity.

### Data pre-processing for model training and inference

We resampled the functional responses and behavior signals (pupil size and locomotion) to 30 Hz—the frame rate of natural movies—using linear spline interpolation. For quality control, we quantified neuronal reliability as the average leave-one-out correlation across repeated stimulus presentation, and the stability as the absolute slope of the linear fit to these correlation values across repeats. Only boutons with reliability greater than 0.4 and stability less than 0.2 were included for model training and further analysis.

To model neural activity as a function of visual stimuli and behavioral state, we appended the behavior signals to the input video, yielding a three-channel model input. We used 80% of the single-repeat movie clips for training and the remaining 20% for validation; movie clips with 20 repeats were reserved for testing. Prior to model fitting, we normalized the videos using the training set to achieve a mean pixel intensity of 0.0 and a standard deviation of 1.0. Behavioral signals and each bouton’s responses were standardized to have a standard deviation of 1.0, and neuronal responses were additionally shifted to be strictly positive on a bouton-by-bouton basis.

### Model training and evaluation

We used Adam optimizer to optimize our network’s parameters (Kingma, 2014). We trained the networks by minimizing the Poisson loss 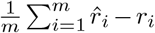, where *m*, 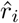 and *r*_*i*_ denote the num-ber of axonal boutons, the predicted response and the recorded response. We did not take into account the natural time ordering for this loss function, i.e., each frame was treated independently. After each training epoch, we computed the correlation between predicted and recorded responses on the validation data, which was averaged across boutons. If the correlation did not increase for five successive epochs, we stopped the training and restored the model to its state with best performance. After that, we either decreased the learning rate by a factor of 0.3 or stopped training altogether, if the number of learning-rate decay steps was equivalent to four.

We used correlation to evaluate predictive performance, which is invariant to shift and scaling of predictions. To measure performance on variation between individual trials, we computed single-trial correlation *ρ*_*st*_ per bouton between predicted responses *o*_*ij*_ and recorded responses *r*_*ij*_ as

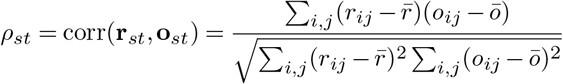

where *r*_*ij*_ represents response to the *i*-th frame of *j*-th video repeat, *o*_*ij*_ represents the corresponding prediction, 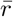 represents the average responses to the videos in test data across all repeats, and *ō* represents the average prediction for the videos in test data across all repeats. We used the averaged of *ρ*_*st*_ across neurons as the final metric.

Similar to *ρ*_*st*_, we computed correlation to average *ρ*_*ta*_ using the average response across video repeats:

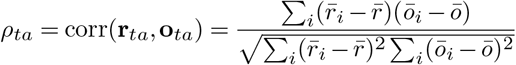

where 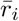 denotes a response averaged over stimulus repeats for one neuron.

### Dynamic most exciting inputs

To generate dynamic MEI for each axonal bouton, we performed gradient ascent on an ensemble of three trained models initialized with different random seeds. We initialized a starting video **x** ∈ ℝ^*h×w×d*^ (stimulus height *h* = 36, width *w* = 64, temporal depth *d* = 40) with Gaussian white noise and fixed the behavior channels to medium values of the training set. In each optimization iteration, we computed the gradients of the response at the last time frame of a single bouton with respect to the input video. To constrain the contrast of the stimulus, we set a L2 norm budget of 50 for the resulting MEI. In total, we ran stochastic gradient descent with a learning rate of 1.0 for 1000 iterations.

### Classifier using readout feature weights

To predict functional types from readout feature weights, we pooled responses to natural movies, chirp and moving bars. In total, the dataset comprised 3,685 functional units from five scans, each with a cell type label assigned by the GMM clustering. Classes containing fewer than 40 boutons were merged into a single dummy label “-1”, resulting in 20 target classes. We trained a multinomial logistic regression classifier using nested cross-validation with 10 stratified folds to predict functional types. Within each training fold, the optimal L2 regularization strength was selected via grid search, and model performance was evaluated on the held-out 10% of data using balanced accuracy. Finally, predictions from the 10 folds were concatenated to construct a confusion matrix, which was normalized such that each row summed to 1.0.

Similarly, we used readout feature weights to predict cortical depth of individual bouton obtained from two-photon structural stack, by pooling 7,005 functional units across five experimental scans and training a logistic regression model with nested cross-validation.

### Parametric analysis of digital-twin predicted responses to looming stimuli

Looming stimuli, generated according to Gabbiani et al. (1999), are parametrized by *l/*|*v*| parameter, where *l* is the half-size of the looming object and |*v*| is the speed of approach. We used looming stimuli with *l/*|*v*| = [20,32,50,80,110] (units: ms).

The generated stimuli were z-scored using the mean and standard deviation of the training video set and presented to the digital-twin to predict individual neuronal responses (number of neurons was 3,685). The responses were grouped according to the chirp and moving bar functional clusters the neurons belong to. Predicted responses were first preprocessed by subtracting a baseline calculated from the initial 3 frames (100 ms) of each response. Then the mean responses of the functional groups were used for further parametric analysis.

To characterize the looming selectivity of functional groups, we applied the parametric model described by Gabbiani et al. (1999). To account for the finite size of the monitor, traces were linearly extrapolated to the theoretical time of physical collision based on the rig geometry (monitor height: 33 cm, platform distance: 15 cm) and the *l*/|*v*| of each stimulus. The time of the peak response relative to collision (*t*_*peak*_) was identified from these mean traces. Then we performed a linear regression of *t*_*peak*_ against the stimulus parameter *l*/|*v*|. The slope of the fitted line, *α*, was used to find the angular threshold of looming object detection: 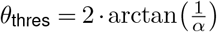.

## Online content

The experimental data is available on Zenodo at https://doi.org/10.5281/zenodo.19041318. The data for training the digital twin model is available on Hugging Face at https://huggingface.co/datasets/open-retina/open-retina/tree/main/franke_lab/qiu_2026. All code to generate results and figures is available on Github at https://github.com/yongrong-qiu/retina-axon-model, as well as the released model weights for *in silico* experiments.

## ACKNOWLEDGEMENTS

This work was supported by the European Research Council (ERC) under the European Union’s Horizon Europe research and innovation programme (grant agreement No. 101117156 to KF, grant agreement No. 101171526 to FS), by the German Research Foundation (DFG; CRC 1233 “Robust Vision: Inference Principles and Neural Mechanisms”, project number 276693517 to FS, KF, TE, PB; CRC 1456, project number 432680300 to FS; ANR/DFG “RetNet4EC”, EU 42/12-1, project number 505379160 to TE, PB), the Hertie Foundation (to PB), the National Science Foundation (grant No. DBI-2021795 to FG, MV; ARNI DBI-2229929 to AST), the National Institutes of Health (grant No. NS130917 to FG, MV; K08NS126573 to CRC, NS128901 to JR), the James Fickel Enigma Project Fund (to AST), and the Intelligence Advanced Research Projects Activity (IARPA) via Department of Interior/Interior Business Center (DoI/IBC; contract number D16PC00003 to AST).

## AUTHOR CONTRIBUTIONS

**Yongrong Qiu**: Conceptualization, Methodology, Software, Formal Analysis, Investigation, Visualization, Writing – Original Draft **Lisa Schmors**: Conceptualization, Methodology, Software, Formal Analysis, Investigation, Visualization, Writing – Original Draft **Na Zhou**: Conceptualization, Methodology, Investigation, Writing – Review & Editing **Mels Akhmetali**: Methodology, Software, Formal Analysis, Visualization, Writing – Original Draft **Dominic Gonschorek**: Methodology, Software, Formal Analysis, Visualization, Writing – Original Draft **Cameron Smith**: Methodology, Software, Investigation, Writing – Review & Editing **Anton Sumser**: Methodology, Investigation **Marie Vallens**: Looming Stimulus Methodology, Writing – Review & Editing **Cathryn Cadwell**: Methodology, Investigation **Fabrizio Gabbiani**: Looming Stimulus Methodology, Writing – Review & Editing **Maximilian Joesch**: Methodology, Investigation **Andreas Tolias**: Resources, Supervision, Funding Acquisition **Philipp Berens**: Resources, Supervision, Funding Acquisition, Writing – Review & Editing **Thomas Euler**: Resources, Supervision, Funding Acquisition, Writing – Review & Editing **Fabian Sinz**: Conceptualization, Supervision, Funding Acquisition, Writing – Review & Editing **Jacob Reimer**: Conceptualization, Supervision, Funding Acquisition, Writing – Review & Editing **Katrin Franke**: Conceptualization, Methodology, Formal Analysis, Investigation, Visualization, Project Administration, Supervision, Funding Acquisition, Writing – Original Draft

## Extended data figures

**Supplemental Fig. S 1.**
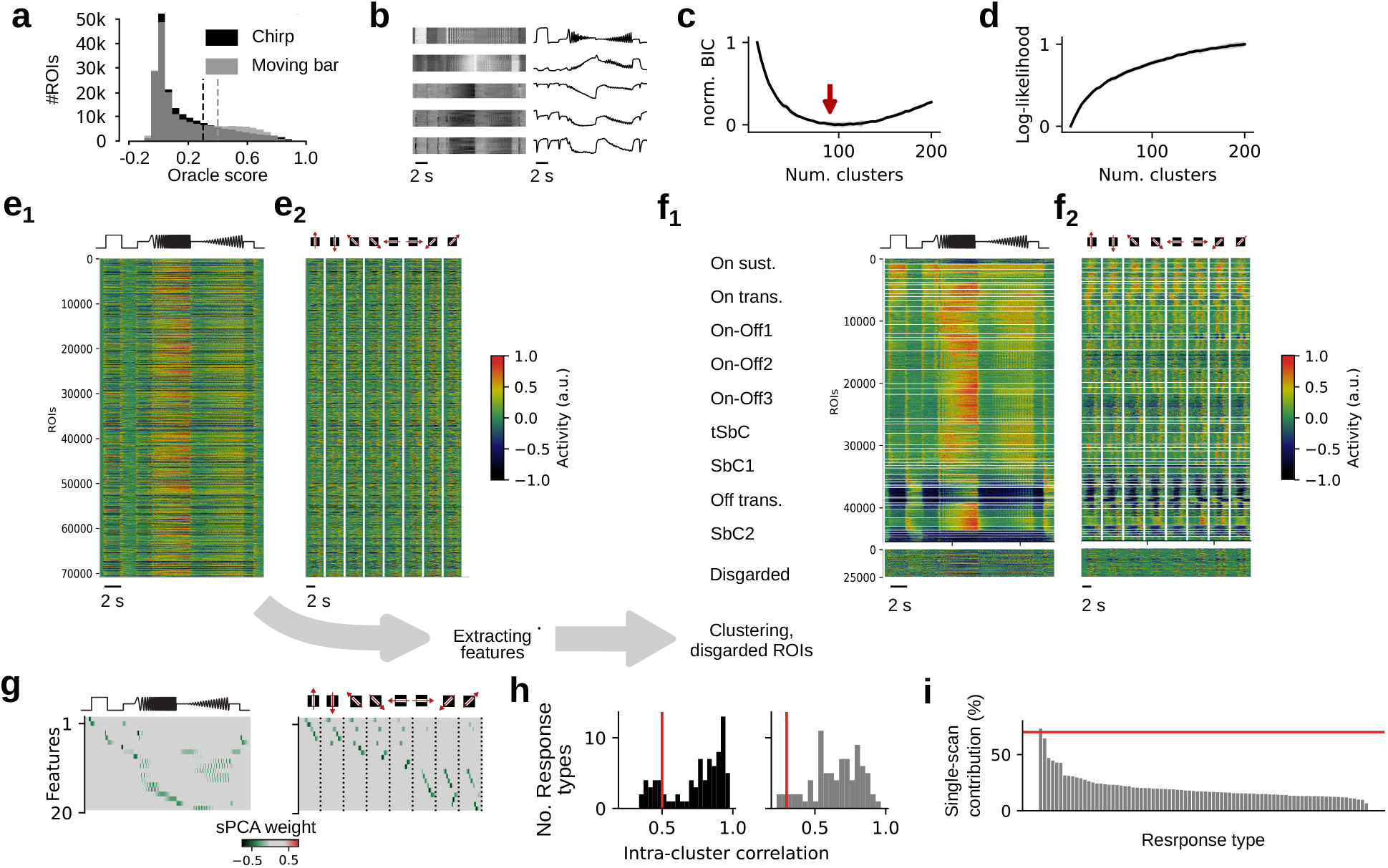
Clustering and data pre-processing. **a**, Signal-to-noise (S/N) ratio of neural responses quantified by oracle correlation (leave-one-out reliability metric) for the chirp (black) and moving bar (gray) stimuli. **b**, Example calcium responses excluded from analysis (left: heat maps; right: mean responses) being artifacts of light-contamination artifacts closeley following stimulus light intensity (top, see (Schmors et al., 2025)) and artifacts due to background-subtraction with sharp negative deflections incompatible with GCaMP off-kinetics (bottom, see Methods). **c**, Normalized Bayesian information criterion (BIC) curve. Arrow indicates the optimal number of clusters. **d**, Normalized log-likelihood of the Gaussian Mixture Model (GMM), saturating with increasing numbers of clusters. **e**, Heat maps of Ca^2+^ responses for (**e**_1_) chirp and (**e**_2_) moving bar of *n* = 71,021 axonal responses before quality filtering. Each row represents the mean response across trials of a single axonal bouton (raw normalized responses). Moving bar responses show all 8 presented directions. **f**, Heat maps of clustered responses (*n* = 50 clusters) for chirp (left) and moving bar (right), including responses discarded based on S/N thresholds and imaging artifacts (bottom, see **a** and **b**). **g**, Temporal features extracted from calcium responses via sPCA (see Methods) and used as input for GMM clustering (**e** and **f**), shown for chirp (left) and moving bar (right). **h–i**, Cluster quality control. **h**, Distributions of intra-cluster correlations computed using a leave-one-out approach. Only clusters with mean intra-cluster correlations above 0.5 (chirp) and 0.3 (bar) were retained to ensure cluster coherence. **i**, Spatial distribution of cluster members across experimental scans, with clusters ranked by their maximum single-scan contribution. Clusters in which a single scan contributed more than 70% of members were excluded as potential batch effects.

**Supplemental Fig. S 2.**
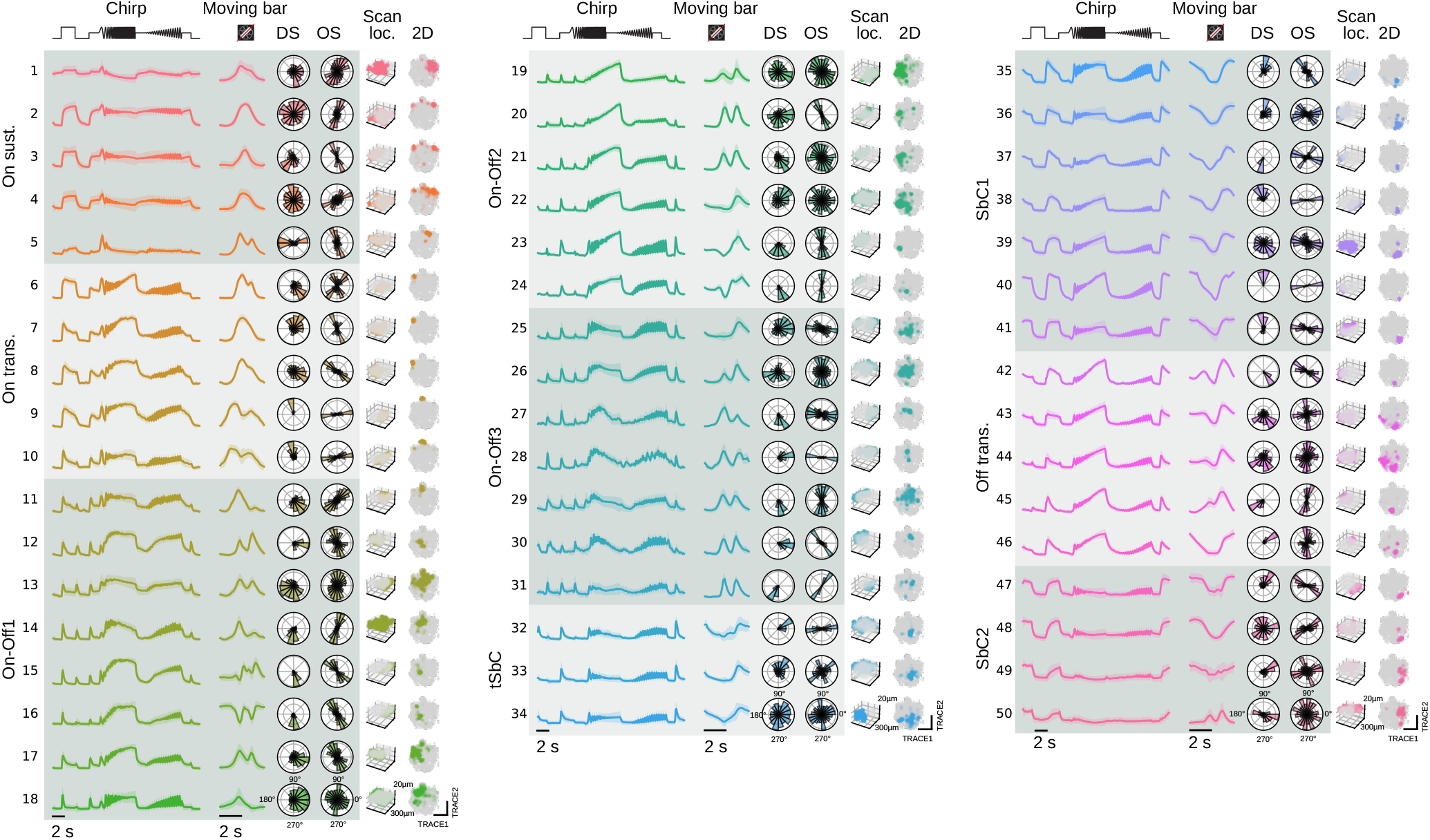
Retinal input modules to SC have diverse functional and anatomical properties. Functional and anatomical characterization of identified clusters with cluster names indicated on the left. Properties from left to right: mean responses to chirp stimulus (shaded area: standard deviation) and moving bar; direction and orientation selectivity with polar plots showing distributions of preferred directions and orientations; anatomical distribution of cluster ROIs within a representative scan field (colored: ROIs of the respective cluster; gray: ROIs not part of the cluster recorded within the same scan field); 2D TRACE (Schmors et al., 2025) embedding showing cluster separation. Anatomical locations show a single scan field (the one with most measured ROIs per cluster) to avoid registration artifacts across scans. (*On sust*.: *On sustained; On trans*.: *On transient; tSbC: transient suppressed by contrast; SbC: suppressed by contrast; Off trans*.: *Off transient*)

**Supplemental Fig. S 3.**
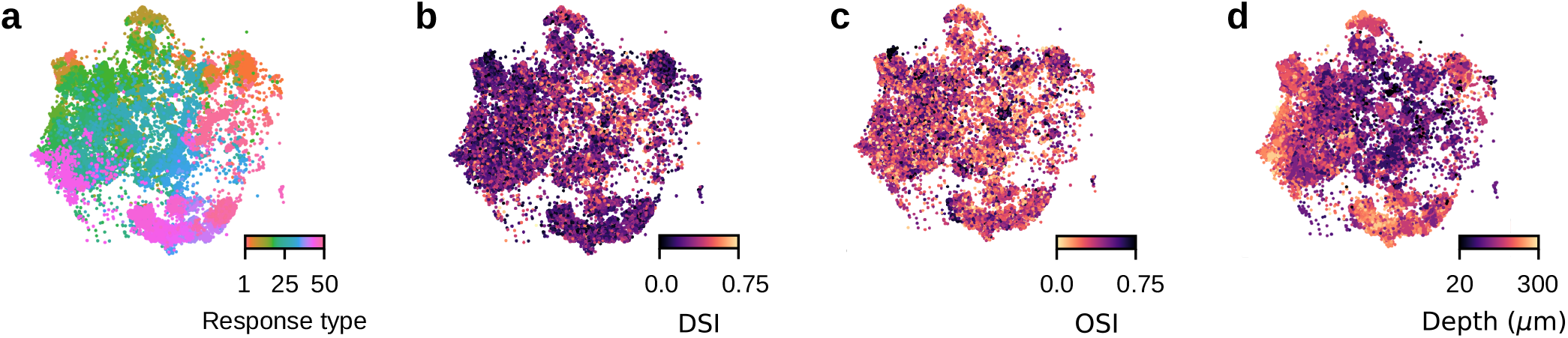
Two-dimensional visualization of response features. Functional responses visualized in 2D using the TRACE algorithm for **a**, response type, **b**, direction selectivity, **b**, orientation selectivity, and **c**, depth,)

**Supplemental Fig. S 4.**
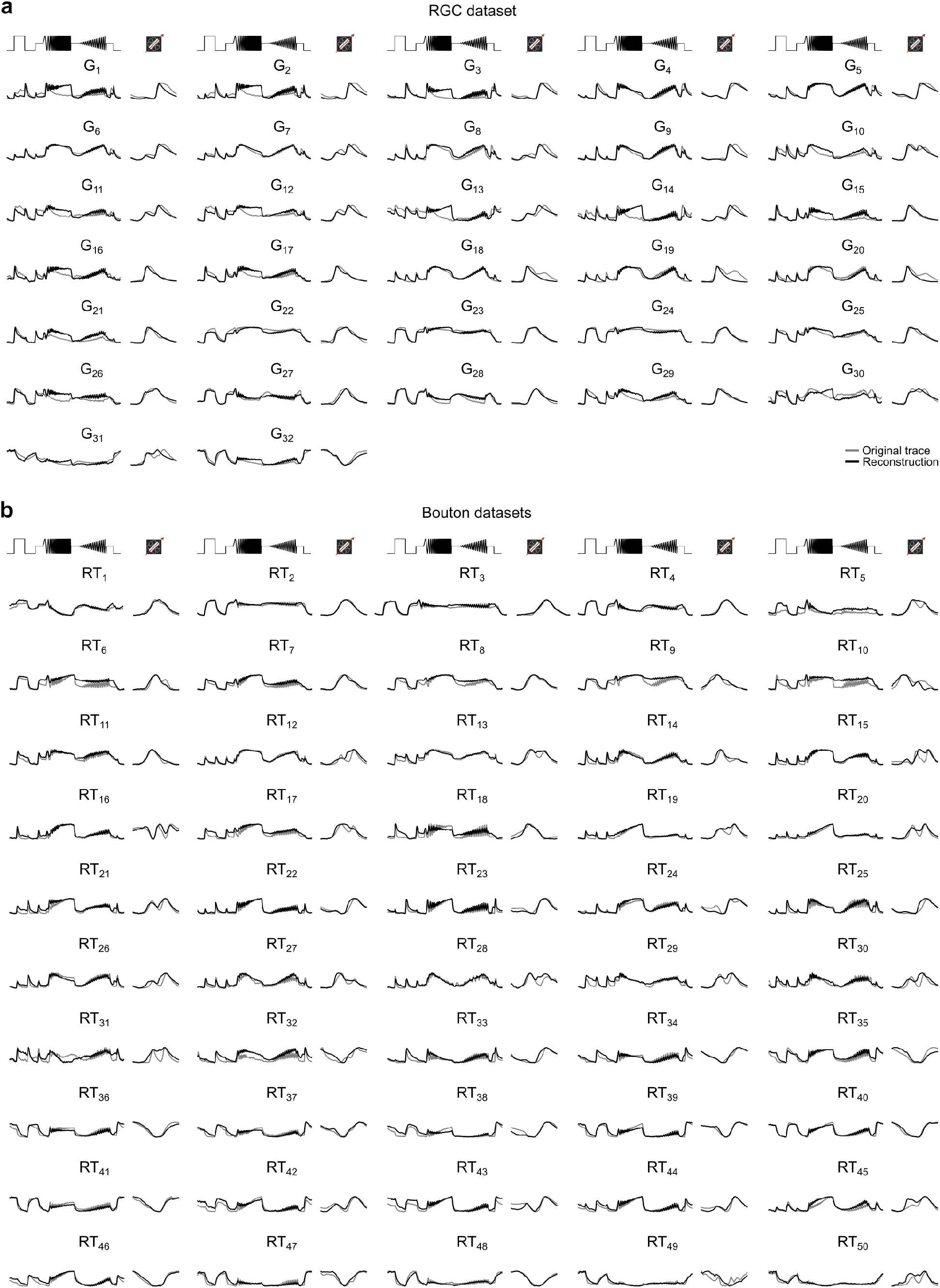
Reconstructed traces of both datasets. **a**, All 32 RGC group responses to the full-field ‘chirp’ (left) and moving bars (right). For each group, mean original traces (gray) and reconstructions (black) are shown. **b**, Same as in a, but for all 50 bouton response types (RT).

**Supplemental Fig. S 5.**
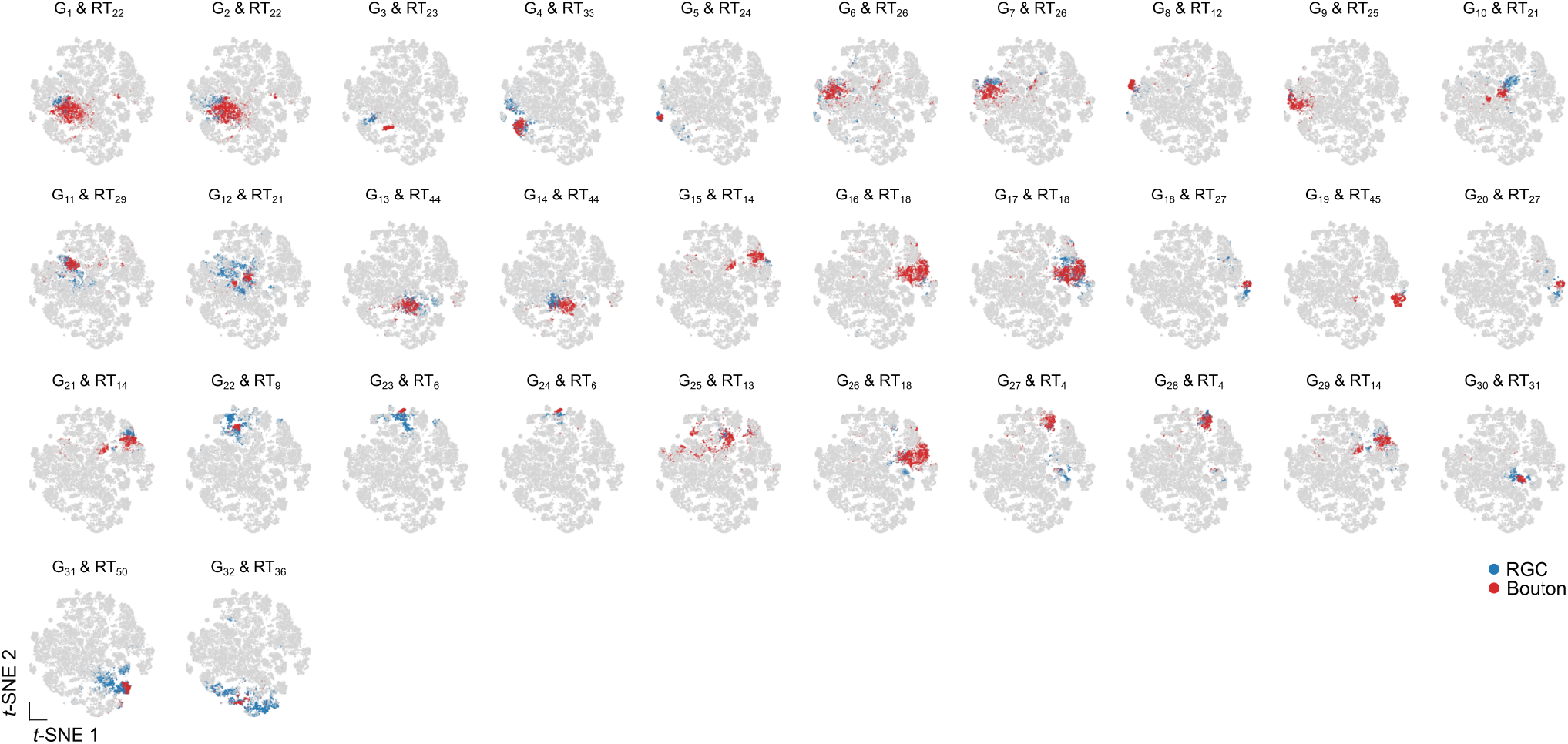
RGC groups mapping to bouton response types. Correspondences between the 32 RGC groups and bouton response types (RT) visualized in the shared latent space visualization using *t*-SNE. Highlighted points show cells from matched groups/types overlaid on the full embedding.

**Supplemental Fig. S 6.**
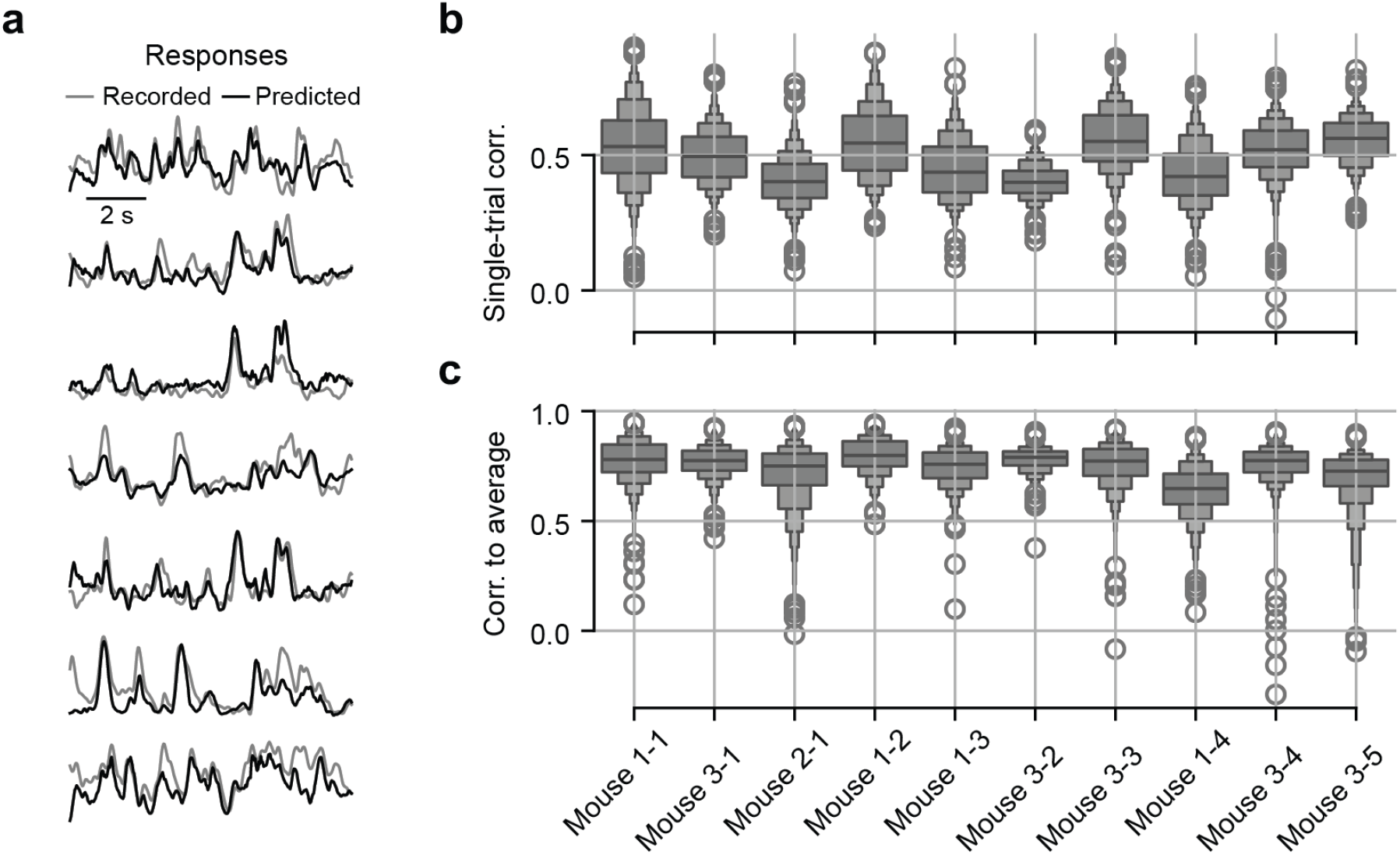
Predictions of our model trained on responses to natural videos. **a**, Example responses of retinal boutons to natural movies (gray, recorded; black, predicted). **b**,**cc** Boxen plots of single-trial correlations (b) and correlations with the trial-averaged responses (c) across 10 scans from three mice, analyzed separately.

**Supplemental Fig. S 7.**
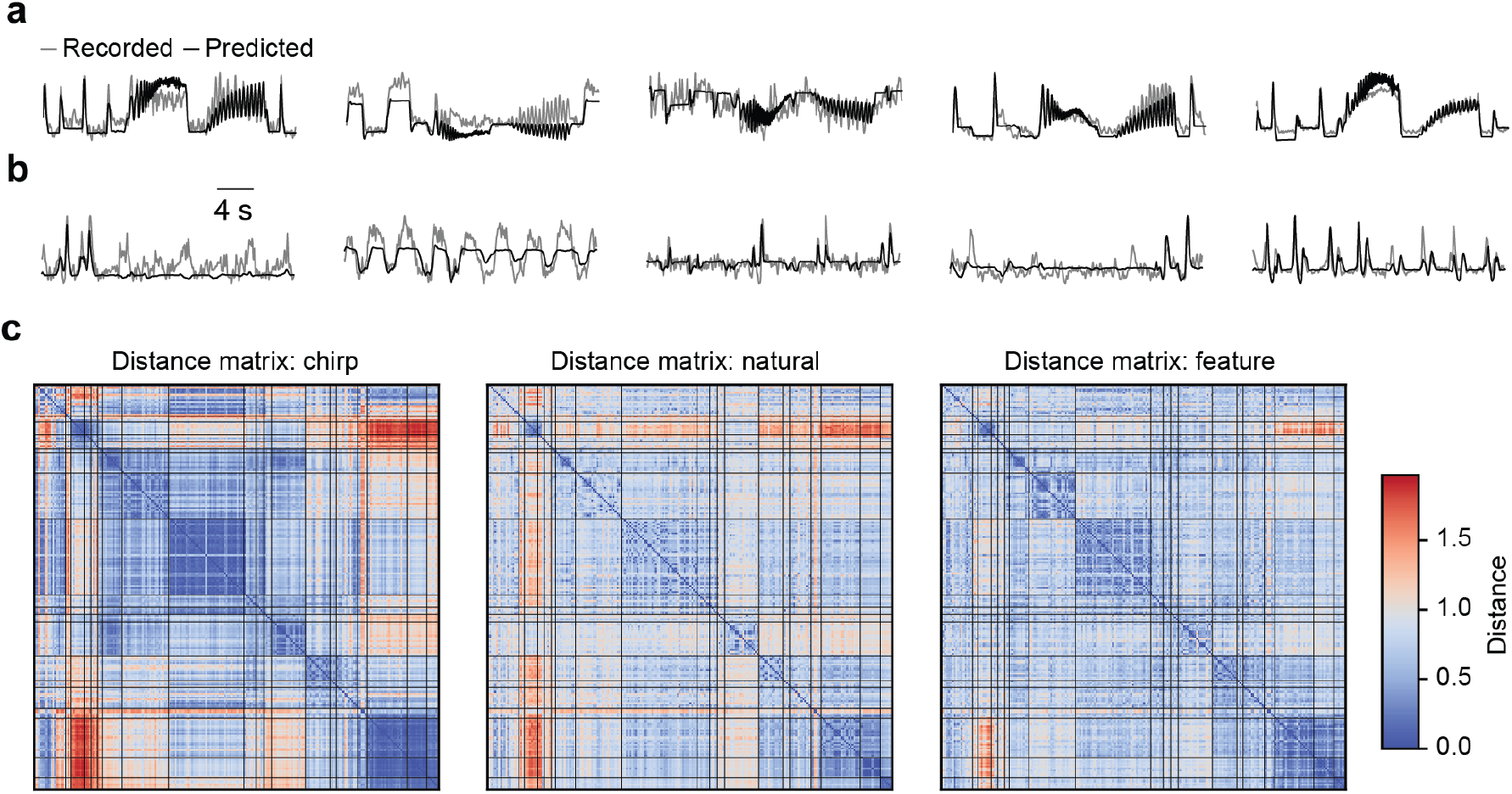
Analysis of predicting response types using readout feature weights. **a**, Responses to the chirp stimulus for exemplary retinal boutons (gray, recorded; black, predicted). **b**, Responses to the moving bar stimulus for exemplary retinal boutons (gray, recorded; black, predicted). **c**, Distance matrices were estimated using responses to chirp stimuli(left), responses to natural movies (middle), and readout feature weights (right) for 3,685 boutons, ordered by their true response types identified via GMM clustering. All three matrices exhibit similar block-diagonal structure, indicating pronounced response diversity across bouton types. To quantify the similarity between these representations, we computed the pairwise rank correlations between distance matrices. The resulting Spearman correlation coefficients were 0.581 for chirp versus natural movies, 0.555 for chirp versus readout features, and 0.626 for natural movies versus readout features. The strong correspondence between the feature-based distance matrix and the response-based matrices suggests that the trained model captures functionally meaningful structure in the neuronal responses.

